# Real-paired single-cell/bulk RNA-seq benchmark and a practical protocol for accurate cell-type deconvolution in human BAL samples

**DOI:** 10.64898/2026.01.14.699304

**Authors:** Yushan Hu, Ziying Liu, Danielle Tsao, Janice M. Leung, Firoozeh V. Gerayeli, Xuan Li, Xiaojian Shao, Don D. Sin, Xuekui Zhang

## Abstract

**Background:** Pseudo-bulk RNA-seq, generated by aggregating single-cell profiles, is widely used for benchmarking deconvolution methods because it globally approximates bulk transcriptomes and provides known cell-type proportions as ground truth. However, pseudo-bulk inherits single-cell specific measurement properties, and the extent to which these differ from real bulk RNA-seq remains difficult to quantify in the absence of paired data. In practice, deconvolution studies also commonly rely on large external single-cell references, yet the value of small, protocol-matched in-study references has not been systematically evaluated under real bulk conditions. These gaps motivate a paired benchmark that jointly examines pseudo-bulk fidelity, reference design, and their consequences for deconvolution accuracy.

**Results:** We establish a fully paired benchmarking framework using split-sample, donor-matched bulk and single-cell RNA-seq (scRNA-seq) from human bronchoalveolar lavage (BAL). Embedded within a reverse five-fold cross-validation design and evaluated on real bulk RNA-seq data, this framework benchmarks 15 deconvolution algorithms published in 2013-2025 across 3 cell-type resolutions and 5 single-cell references, including three published BAL datasets, a harmonized lung BAL atlas, and an in-study reference derived from paired aliquots. We show that real bulk and matched pseudo-bulk profiles exhibit systematic gene-level differences, identifying 557 reproducibly discordant genes (|log_2_ FC| *>* 1, FDR*<* 0.05 by LIMMA), including cell-type informative features. These discrepancies reflect technology-specific effects and violate the linear mixing assumption underlying most deconvolution methods. We demonstrate that a protocol-matched in-study reference constructed from only six donors consistently outperforms substantially larger external references, with the advantage that increases at finer cell-type resolution. Moreover, paired samples enable the identification and selective removal of discordant genes, which further improves deconvolution accuracy for many algorithms, particularly in high-resolution settings. These findings are robust across methods, references, and evaluation criteria and extend beyond compositional accuracy to improving recovery of disease-associated cell-type differences in clinical applications.

**Conclusions:** Our study provides the first fully paired benchmark of transcriptomic deconvolution on real bulk RNA-seq data of human BAL samples and demonstrates that reference design and data compatibility are as influential as algorithm choice. Beyond benchmarking, we introduce a practical and cost-effective protocol for deconvolution studies: generate single-cell data for a minimal subset of bulk samples (pilot pairing), use these data to construct an in-study reference and identify discordant genes, and apply the resulting insights to the full cohort. This strategy requires limited additional experimental effort yet yields substantial gains in accuracy and stability, offering actionable guidance for future bulk RNA-seq deconvolution studies across tissues and platforms.

**One-sentence summary:** Systematic benchmarking reveals pseudo-bulk biases and provides practical fixes for accurate RNA deconvolution.

## 1 Background

Inferring cell-type composition from transcriptomic data has become indispensable because virtually every complex tissue, from rapidly evolving tumors to chronically inflamed organs and developing embryos, behaves as a dynamic mosaic of interacting cell populations whose frequencies shift with genotype, disease course, and therapy [1, 2, 3]. Knowledge of those shifts enables mechanistic insight, biomarker discovery, and rational intervention across diverse fields. In oncology, for instance, quantifying immune-cell infiltration and stromal heterogeneity helps predict and monitor responses to checkpoint blockade and other immunotherapies [4, 5]; in neuroscience, charting neuronal, glial, and vascular proportions illuminates region-specific vulnerability in neurodegenerative disorders [4, 6]; and in autoimmunity or infection, resolving the balance between effector and regulatory subsets reveals the cellular circuits that drive pathology [7, 8].

Single-cell RNA-seq (scRNA-seq) provides an unparalleled, cell-type-resolved view of the transcriptome and excels at discovering rare or previously unrecognized phenotypes [9]. Yet generating thousands of high-quality single-cell libraries remains extremely costly, labour-intensive for large cohorts and subject to large noise-to-signal variation owing to multiple factors including batch effect, is constrained by limited sequencing depth, and can introduce artificial shifts in gene expression during tissue dissociation and capture [10]. Bulk RNA-seq, in contrast, is relatively inexpensive, robust and already available for tens of thousands of clinical and population-scale samples, but its aggregate signal masks cell-type-specific biology. *In-silico* deconvolution closes this gap. By leveraging reference profiles often derived from scRNA-seq, it infers the cellular composition of bulk libraries without further experimentation, thereby circumventing dissociation bias and enabling composition studies in very large or otherwise intractable cohorts [11]. These inferred proportions now guide translational decisions ranging from tumor-immune stratification prior to checkpoint blockade to retrospective mining of archived clinical trials, amplifying the need for rigorous benchmarks and transparent experimental guidelines.

Transcriptomic deconvolution formulates the bulk expression vector *b* ∈ ℝ*^G^* as a linear (or sometimes probabilistic) mixture of cell-type signatures *R* ∈ ℝ*^G^*^×*K*^ weighted by unknown proportions *w* ∈ ℝ*^K^*:

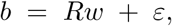

where *G* is the number of genes, *K* the number of cell types, and *ε* captures noise and model mismatch. Algorithms seek either (i) *ŵ* alone or (ii) both *ŵ* and R^ under additional constraints [12, 2, 13, 14]. Approaches span non-negative least squares [12], support-vector regression with noise modelling [2], multi-subject weighted regression [13], Bayesian latent-variable models [14] and multi-objective gene-selection frameworks that optimize for cell-type separability [15]. Adoption is widespread: digital cytometry now accompanies most the Cancer Genome Atlas (TCGA) pan-cancer studies, deconvolution of brain banks anchors neurodegenerative meta-analyses, and immunology consortia routinely integrate single-cell references with bulk cohorts to map immune trajectories.

Current benchmark studies adopt a variety of “ground-truth” strategies, each revealing different aspects of deconvolution performance but none providing a fully paired bulksingle-cell test dataset. When true bulk RNA-seq data are available, validation often relies on an independent measurement method. For immune-rich tissues, flow cytometry (FACS) has been the preferred choice; e.g. Sturm *et al.* quantified peripheral blood mononuclear cells (PBMC) subsets to reveal method-specific pitfalls [4]. Additionally, FACS requires cell dissociation and large sample volumes and is limited by marker availability. Immunohistochemistry (IHC) and RNA Scope offer spatial resolution within tissue sections, preserving structural context; however, like FACS, they depend on specific markers and may not fully capture all cell types, particularly in heterogeneous tissues [16, 6].

The majority of large-scale benchmarks build pseudo-bulk directly from scRNA-seq. Jin & Liu injected controlled noise to probe robustness [7], and Avila Cobos *et al.* showed that preprocessing approaches can rival algorithm choices in impact [8]. Such simulations offer precise control over composition but inevitably miss the full complexity and protocol bias of *in vivo* tissue. Hybrid designs seek to balance realism and control: Tran *et al.* combined breast cancer bulk data with scRNA-derived pseudo-bulk [5], and Xu *et al.* produced guideline-oriented evaluations across mixed scenarios [17]. However, mismatched structure between synthetic and real samples can blur conclusions.

Only a handful of studies approximate a paired design. Sutton *et al.* generated bulk, nuclear, and snRNA-seq data from the same frozen brain blocks (n = 5), but the three modalities were taken from adjacent rather than identical aliquots and the results may have been confounded by nucleus-specific expression biases. Dai *et al.* leveraged bulk, snRNA-seq, and IHC from ROSMAP and other cohorts, but the datasets came from different individuals and across different brain regions [18]. Cobos *et al.* (2023) reviewed *in vitro* mixtures and disparate public bulk datasets without any true pairing [19]. Huuki-Myers *et al.* integrated bulk cortical RNA-seq with orthogonal assays, yet sample size was small and tissue was restricted [6].

Collectively these efforts have advanced the field, but they still hinge on simulated mixtures, approximated pairings, or limited marker panels, leaving open several critical questions. For example, how do algorithms perform when confronted with **real, perfectly matched** bulk and single-cell seq data? Or how many paired samples are necessary, which genes require filtering, and how does annotation granularity affect the accuracy of deconvolution?

To address these and other limitations, we designed a split-sample study of human BAL and performed the first fully paired benchmark. Thirty human BAL samples were divided at collection: one aliquot generated a bulk seq library, the other a 10x Chromium scRNA-seq library (median 5,384 cells). We benchmark 15 deconvolution algorithms across 15 scenarios, combining five single cell reference datasets with three annotation resolutions. The references comprise three published BAL datasets, a harmonized lung BAL atlas, and an in-study reference from 30 paired aliquots. The annotation resolutions are 11 lineages, 13 intermediate types, and 18 fine sub-clusters, with the latter two reflecting progressively finer subdivisions of macrophage states, which are by far the most abundant immune cells in human BAL. Differential analysis between true bulk and paired pseudo-bulk reveals 557 discordant genes (| log_2_ FC| *>* 1, FDR *<* 0.05, by LIMMA), elucidating protocol-driven violations of the linear-mixing assumption. Filtering these genes improves most methods by 18 percentage points in *R*^2^ performance, with effectiveness varying by cell type resolution (finer resolutions show greater benefit: Level 3 vs. Level 1, p<0.001) and filtering method (LIMMA, edgeR, DESeq2, advanced DE methods outperform simple paired *t*-tests, p<0.01) [20, 21, 22]. We translate these insights into a three-step protocol (pilot pairing, in-study reference, and discordant-gene filter), and we release processed data at GSE.

## 2 Results

### 2.1 Dataset overview

To create a benchmarking setting that mirrors real-world bulk RNA-seq deconvolution, we generated a fully paired dataset in which each donor contributed one aliquot for bulk RNA-seq and one for scRNA-seq. We processed in-study lung BAL samples from 32 donors (4 from stable parients with chronic obstructive pulmonary disease, COPD; and the rest from volunteer participants’ without any known chronic lung condition). After quality control, two non-COPD scRNA-seq samples were removed, yielding 241,924 high-quality single cells. These cells were annotated at three resolutions: 11 major types, 13 intermediate types, and 18 fine-grained macrophage-related sub-clusters, capturing lineage structure and macrophage-state heterogeneity (Fig. 2) [23]. Cell-type abundances varied across donors, with some rare subsets absent in individual samples; full cell-type distributions are shown in Fig. S5.

**Figure 1:**
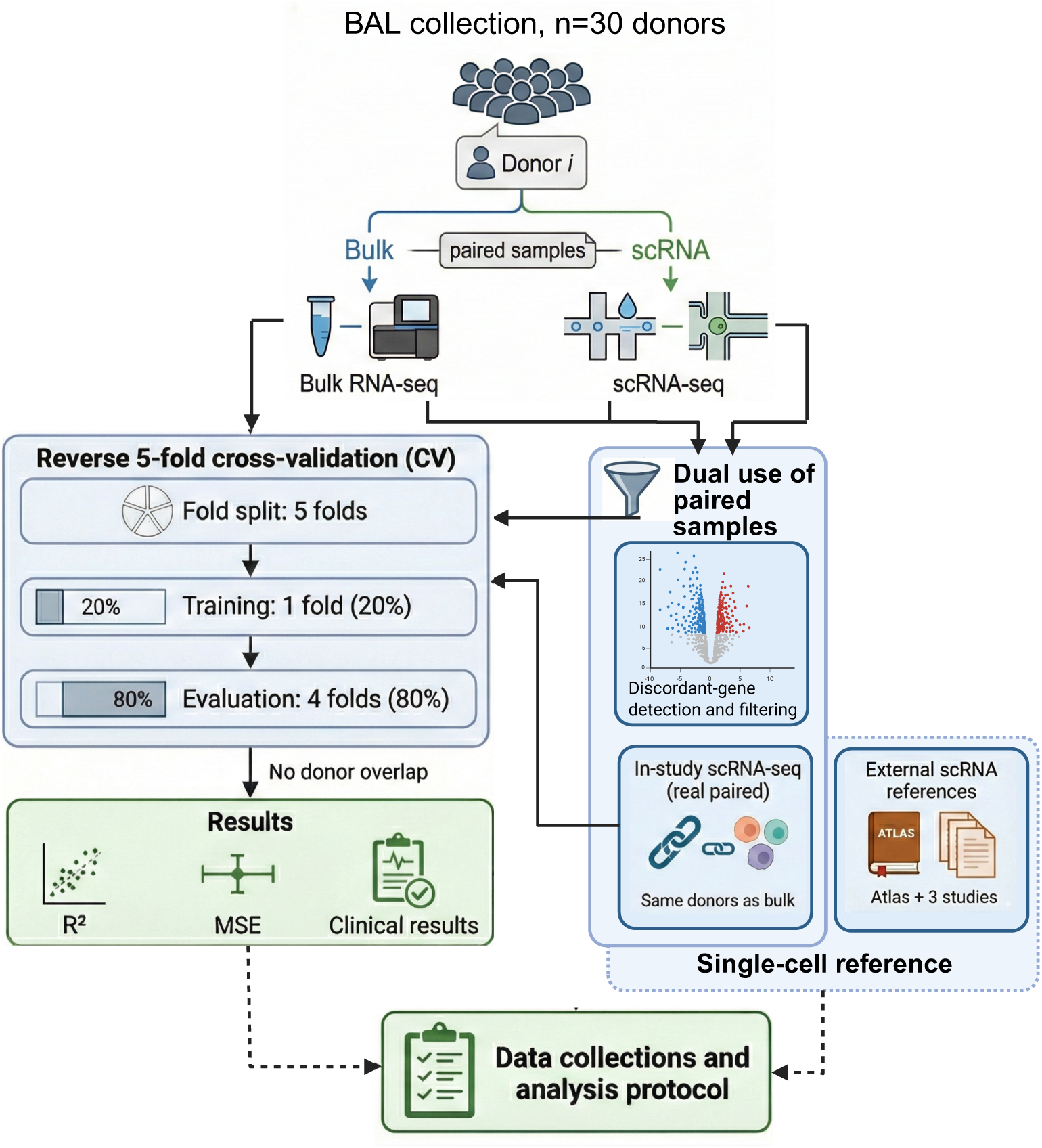
Study design and cross-validation framework using paired bulk and single-cell RNA-seq data from the same BAL sample. Split samples from 30 BAL donors were processed for both bulk RNA-seq and scRNA-seq. Single-cell data were used to construct cell-type references (either from the same donors or external studies), while paired data were subjected to discordant gene filtering. Deconvolution methods were evaluated using reverse 5-fold cross-validation with 20% training and 80% evaluation sets, ensuring no donor overlap between folds. Performance was assessed using *R*^2^, mean squared error (MSE) and clinical outcome predictions.

**Figure 2:**
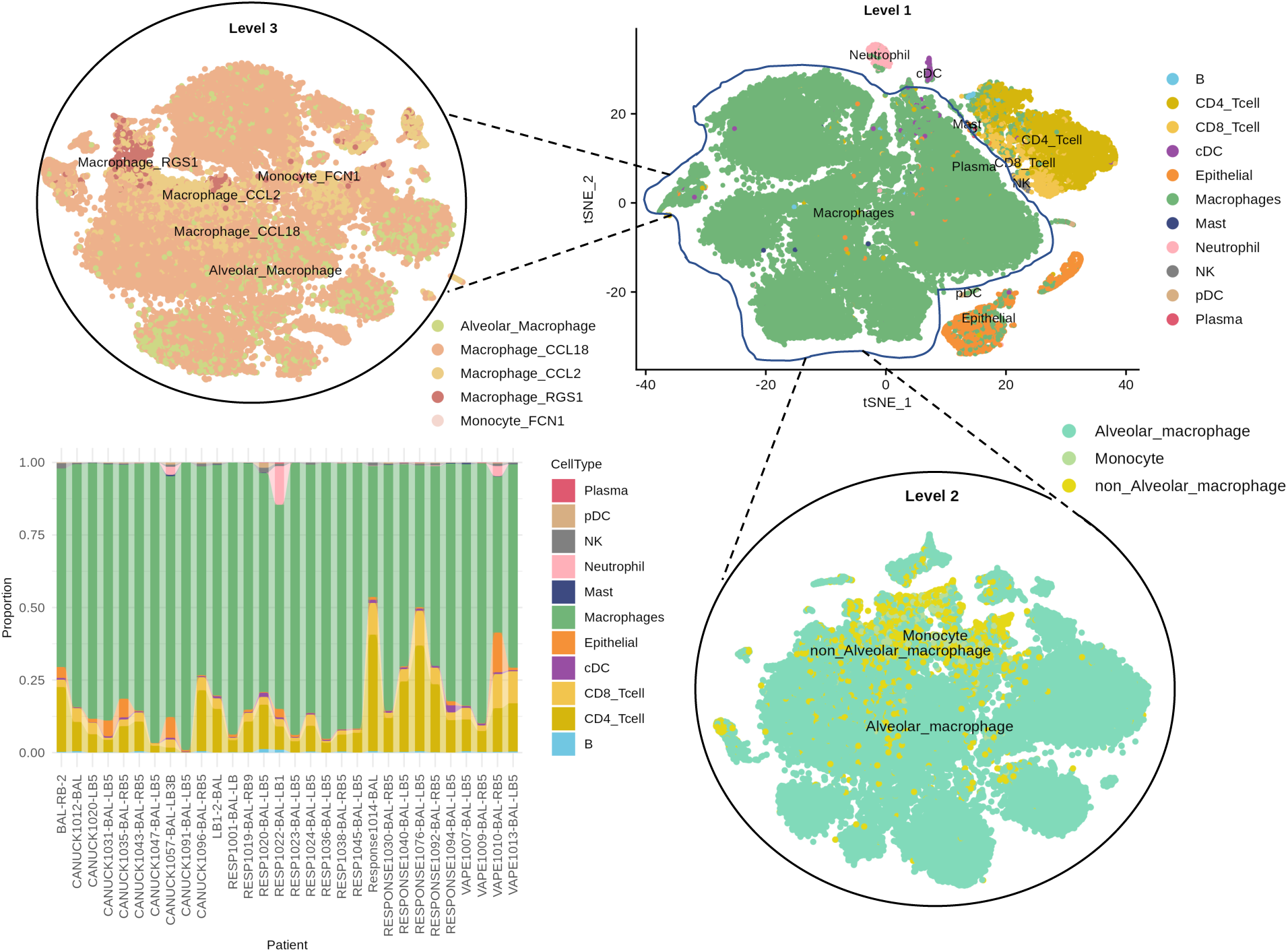
UMAP of in-study single-cell data with three cell-type resolutions and stacked bar plots of cell-type proportions. Level 1: Eleven broad cell types include B cells, CD4^+^ T cells, CD8^+^ T cells, conventional dendritic cells, epithelial cells, macrophages, neutrophils, mast cells, NK cells, plasmacytoid dendritic cells, and plasma cells. Level 2: Thirteen intermediate types a subdivision of macrophages (alveolar macrophage, non-alveolar macrophage, monocyte), resulting in thirteen categories. Level 3: Eighteen fine sub-clusters representing a further division of non-alveolar macrophage states (CCL18^+^, CCL2^+^, RGS1^+^, MT1G^+^) and monocyte subsets (FCN1^+^, IL1B^+^).

A key feature of our evaluation framework is a *reverse* five-fold cross-validation scheme. In each fold, six paired donors (20% of the cohort), not including patient samples, are used to build an `in-study` scRNA-seq reference, while the remaining 24 donors serve as evaluation samples. This design has two benefits: (i) it strictly avoids information leakage because no donor’s scRNA-seq profile ever appears in both the reference and the ground truth; and (ii) it directly tests how well deconvolution performs when only a *small* number of paired donors are available, a common constraint in practice. Demonstrating that six-reference-donor signatures can outperform much larger external references provides evidence that modest pilot pairing is feasible and advantageous.

To systematically assess how biological and technical compatibility of reference panels influences deconvolution performance, we additionally incorporated four publicly available BAL scRNA-seq datasets (Table 1) as alternative references. These datasets were harmonized and, when necessary, restricted to healthy individuals to ensure comparability. By combining in-study and external scRNA-seq references, we are able to directly evaluate how protocol concordance, sample origin, and annotation granularity impact deconvolution accuracy.

**Table 1:** BAL datasets used in this project.

| Study | Data type | Sample size | Cells Count | Genes Count |
| --- | --- | --- | --- | --- |
| In-house | Bulk RNA-seq | 30 | – | 63,187 |
|  | scRNA-seq | 30 | 241,924 | 33,465 |
| Wauters | scRNA-seq | 35 | 65,166 | 33,538 |
| Altas | scRNA-seq | 26 | 352,732 | 63,953 |
| Li | scRNA-seq | 7 | 113,213 | 27,805 |
| Liao | scRNA-seq | 13 | 66,452 | 23,916 |

### 2.2 Real vs. simulated bulk matters

Pseudo-bulk aggregation is widely used for benchmarking deconvolution methods and has produced many influential studies. However, because pseudo-bulk inherits properties of single-cell assays, its similarity to true bulk can only be quantified when real paired samples are available. Our design allows this comparison directly.

To quantify the concordance between real bulk RNA-seq and matched pseudo-bulk profiles, we performed differential expression (DE) analysis using all 30 BAL samples, including both healthy controls and COPD cases. Bulk RNA-seq contained 63,187 genes, and pseudo-bulk profiles were generated by aggregating single-cell expression within each donor (13,915 genes). After taking the gene intersection, 11,397 common genes were retained for paired comparison.

Real bulk RNA-seq data (transcripts per million (TPM)-normalized) and pseudo-bulk data aggregated from scRNA-seq (counts per million (CPM)-normalized) were Z-score normalized across samples to enable a fair comparison of two technologies on a similar scale. As a sensitivity analysis, we also performed *log*_2_-transformation prior to Z-score normalization. DE analysis was performed using LIMMA with empirical Bayes moderation, treating each donor as a paired observation. Across the cohort, we identified 557 discordant genes (≈ 4.9% of the 11,397 tested) showing significant differences between real and pseudo-bulk expression (|log_2_ FC| *>* 1, FDR *<* 0.05), as illustrated in Fig. 3**A**. To assess the robustness of these findings, we additionally repeated the analysis using paired *t*-tests, DESeq2, and edgeR. All four approaches yielded broadly consistent sets of discordant genes (Supplementary Fig. S2), confirming that the observed differences reflect reproducible platform-specific effects rather than artifacts of a particular statistical method.

**Figure 3:**
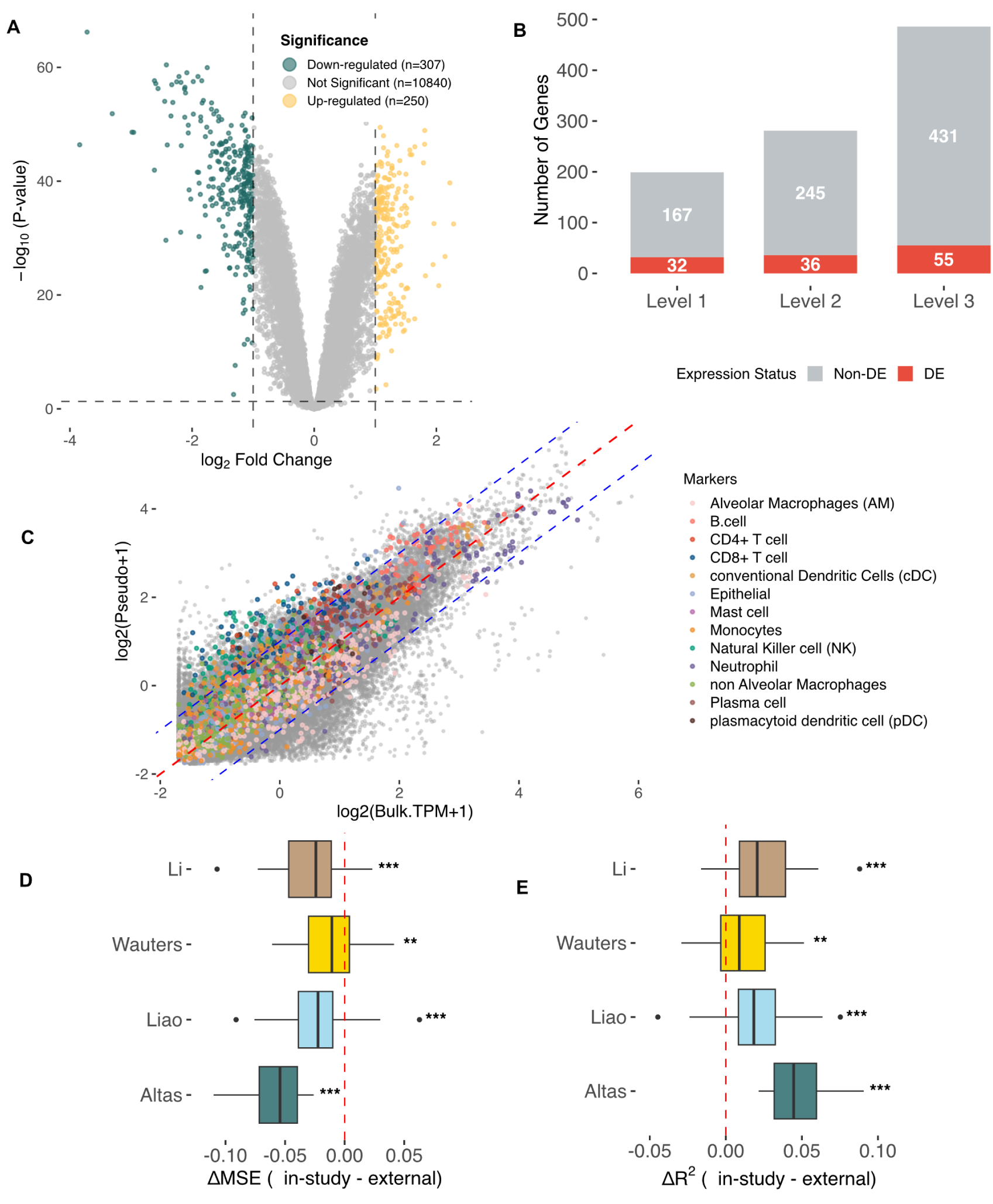
Comparison between real and pseudo-bulk transcriptomic profiles. (**A**) Differential expression (LIMMA) between real and pseudo-bulk RNA-seq. (**B**) DE vs Non-DE across feature genes used to distinguish cell-type. (**C**) Gene-wise scatter plot of log_2_ expression in real vs. pseudo-bulk. (**D–E**) Fidelity of pseudo-bulk generated using alternative single-cell references, evaluated against true bulk across 30 donors (Wilcoxon signed-rank test; \*\*\**p <* 0.001, \*\**p <* 0.01, \**p* < 0.05).

Although the vast majority of genes are concordant, the discordant subset includes canonical marker genes and high-loading features commonly used in signature matrices (Fig. 3**B**). These deviations are consistent with well-known protocol characteristics, such as polyA^+^ fragmentation in bulk versus 3’-end Unique Molecular Identifier (UMI) capture in single-cell assays, gene-length sensitivity, and other biochemical factorsthat introduce small but reproducible technology-driven shifts in expression, leading to systematic deviations from the linear-mixing assumption *b* = *Rw*. Such shifts, while subtle in magnitude, directly affect genes that disproportionately influence deconvolution models and, therefore, warrant explicit quantification in paired bulk-single-cell datasets. Figure 3**C** summarizes overall concordance: real and pseudo-bulk profiles correlate strongly (*r* = 0.853, *R*^2^ = 0.728), with 93.9% of genes falling within ±1-fold difference. This confirms that although the discordant subset is a minority of genes, it involves a lot of cell-type marker genes that disproportionately influence deconvolution.

While discordance between real bulk and pseudo-bulk generated from paired single-cell profiles primarily reflects technology-specific differences, the use of external single-cell references can introduce additional deviations from the linear-mixing assumption (*b* = *Rw*) underlying deconvolution. To quantify this effect, we constructed ‘reference-derived’ bulk profiles for each donor by multiplying each scRNA-seq reference matrix *R* with the donor-specific true cell-type proportions *w* estimated from our paired single-cell data. For each reference and donor, we then computed the mean squared error (MSE) and coefficient of determination (*R*^2^) between the reference-derived bulk and the observed bulk RNA-seq.

To compare the in-study reference with the four external references on a common scale, we summarized performance using pairwise differences

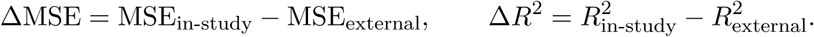

Negative values of ΔMSE therefore indicate that the `in-study` reference yields *lower* error (i.e. external references introduce more deviation from the true bulk), whereas positive values of Δ*R*^2^ indicate that the `in-study` reference explains *more* variance than the external reference. As shown in Fig. 3**D–E**, the distributions of ΔMSE are predominantly below zero and those of Δ*R*^2^ above zero across the 30 donors for all four external references, and these shifts are statistically significant (Wilcoxon signed-rank test, *p <* 0.01). The systematic pattern in Fig. 3**D–E** indicates that a protocol-matched `in-study` reference yields bulk reconstructions that more closely resemble true bulk transcriptomes than those derived from larger external datasets. Because many deconvolution algorithms depend directly on the linear-mixing model *b* = *Rw*, such improvements in fidelity have two consequences. First, they imply that the `in-study` reference provides a more accurate basis for constructing the signature matrix *R*, reducing technology-driven discrepancies in cell-type defining genes. Second, they suggest that downstream evaluation is also more reliable when the reference and bulk profiles share the same biochemical and library-generation features. Together, these observations support the hypothesis that even a relatively small number of paired donors may outperform substantially larger external references for most deconvolution methods. This motivates the analyses in Section 2.4, where we formally quantify the extent to which an `in-study` reference confers performance gains across various deconvolution methods and three annotation resolutions.

### 2.3 Deconvolution performance across methods, references and annotation resolutions

We assessed 15 deconvolution algorithms across three hierarchical annotation levels (11 lineages, 13 intermediate types, and 18 macrophage-resolved sub-clusters) and five single-cell reference datasets, generating a comprehensive landscape of how algorithm choice interacts with reference composition and cell-type granularity. Together, the four panels in Fig. 4 summarize inter-method performance from Level 1 to Level 3 (Fig. 4**A–C**), cross-resolution behavior from Level 1 to Level 3 (Fig. 4**D**), and reference-driven variability across all levels (Fig. 4**E–G**).

**Figure 4:**
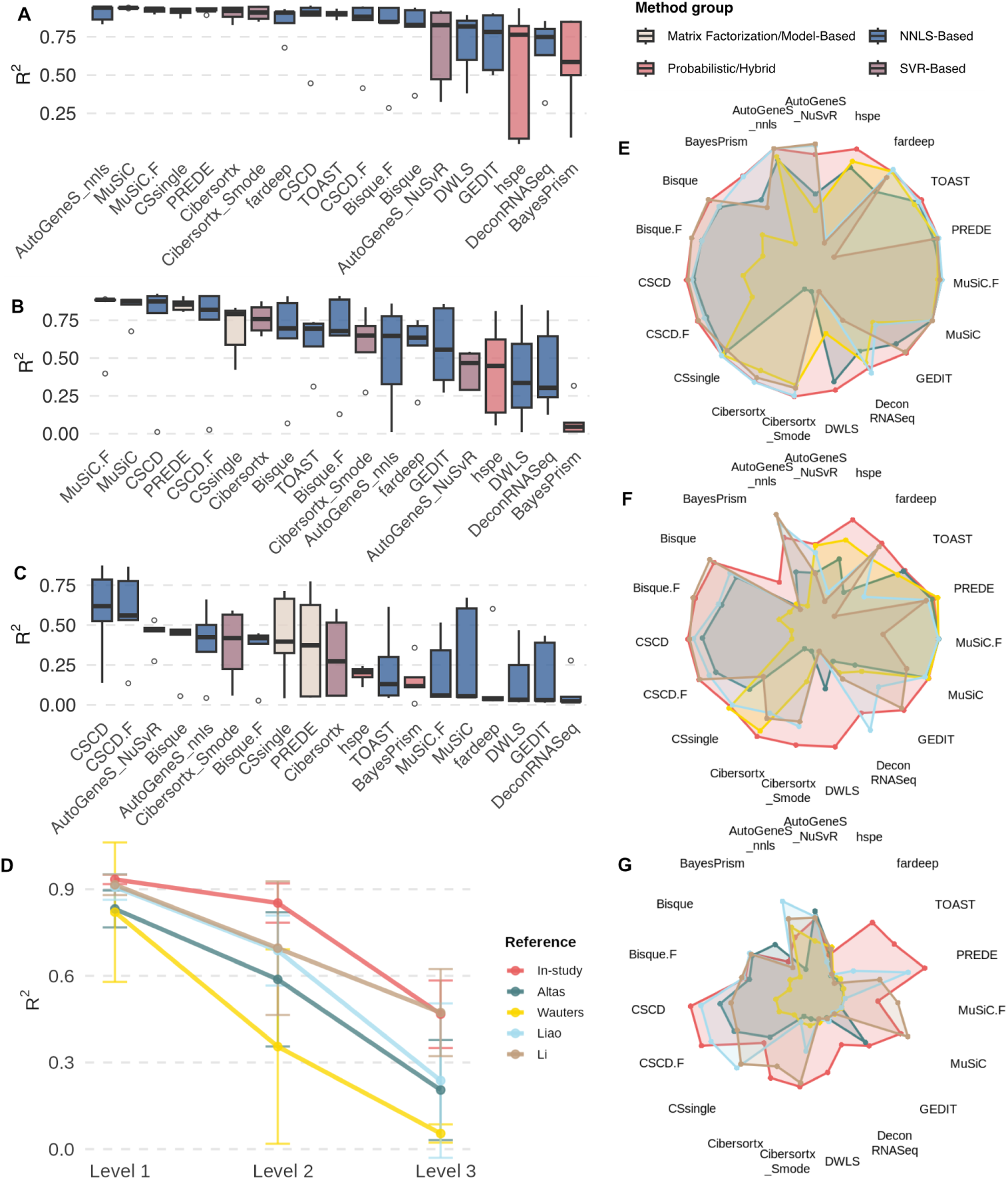
Comprehensive evaluation of deconvolution method performance across three levels and five references. (**A—C**) *R*^2^ performance by the deconvolution method from Level 1 to Level 3. Boxplots show distribution across reference datasets, colored by method group. Methods ordered by median performance. (**D**) Cross-resolution performance comparison. Boxplots show *R*^2^ distributions across alternative references (gray/beige/green for Levels 1/2/3). (**E—G**) Radar charts comparing method performance across five reference datasets at Level 1, 2, and 3, respectively. Colored lines represent references: `in-study` (pink), BAL atlas (teal), Liao *et al.* (blue), Wauters *et al.* (yellow), Li *et al.* (brown). Radial distance indicates *R*^2^ (0-1). Polygon size decreases with finer resolution.

At the intermediate resolution of 13 cell types, which provides a practical balance between biological interpretability and signature complexity, NNLS-based and matrix factorization/model based approaches achieved the strongest performance when paired with the in-study reference (Fig. 4**a,b**). CSCD, CSCD.F, and MuSiC.F reached median *R*^2^ values exceeding 0.95, reflecting the advantage of linear models with shrinkage or adaptive weighting when expression signatures are closely matched to the bulk cohort. PREDE and CSsingle also performed competitively (group median *R*^2^ ≈ 0.93), indicating that non-negative factorization can be robust under modest reference mismatch. In contrast, both SVR-based methods (CIBERSORTx and AutoGeneS) and probabilistic/hybrid models (BayesPrism and HSPE) displayed broader performance dispersion, indicating greater sensitivity to gene-level noise and inter-study variability. MSE metrics mirrored these findings: NNLS-based methods yielded the lowest reconstruction error, matrix factorization/model based methods followed closely, whereas several SVR and probabilistic approaches exhibited substantially higher error ranges (MSE distributions shown in Supplementary Fig. S1).

Increasing annotation resolution markedly reduced absolute performance across all methods and references, but the magnitude of this decline depended strongly on both the reference and the algorithm (Fig. 4**D**). From Level 1 (11 broad lineages) to Level 3 (18 macrophage and monocyte sub-clusters), median *R*^2^ values dropped from the 0.6–0.9 range to near zero for many method reference combinations, consistent with the expected increase in signature overlap and multicollinearity at finer granularity. Notably, the `in-study` reference exhibited a substantially slower decay in *R*^2^ than the external references, maintaining moderate accuracy even at Level 3. This behavior indicates that protocol-matched references can partially offset the increased difficulty of subtype-level deconvolution. A subset of algorithms, including CSCD, CSCD.F, Bisque, Bisque.F, CSsingle, and AutoGeneS, remained comparatively robust: although their *R*^2^ values decreased with resolution, they still retained median *R*^2^ ≈ 0.4–0.6 at Level 3 (Fig. 4**C**), suggesting that stronger regularization, adaptive gene weighting, or internal feature optimization can mitigate some of the complexity introduced by fine-grained annotations.

Changing the single-cell reference had a profound effect on both absolute performance and method rankings, as visualized by the radar plots in Fig. 4**E—G**. With the `in-study` reference (pink polygons), most methods achieved large, relatively symmetric profiles at each level, indicating high *R*^2^ and comparatively balanced performance across cell types. In contrast, polygons corresponding to external references (`In-study`, BAL atlas, Liao *et al.*, Wauters *et al.*, Li *et al.*; pink, teal, blue, yellow, and brown, respectively) were smaller and often skewed, reflecting both reduced accuracy and uneven performance across cell types. These differences were particularly pronounced at Levels 2 and 3 (Fig. 4**F,G**), where some probabilistic and hybrid methods collapsed to very small polygons, consistent with near-zero *R*^2^ observed for the Li *et al.* reference. Importantly, the identity of the “best” method was not stable across references: while NNLS- and SVR-based methods dominated when using the `in-study` reference, matrix factorization/model-based approaches (notably PREDE) became top-ranked with the Wauters reference, and the relative order of SVR versus NNLS methods inverted for some external datasets. MuSiC (NNLS family) and PREDE (matrix-factorization family) were among the most stable methods at coarse and intermediate resolutions, whereas HSPE and BayesPrism exhibited pronounced volatility across nearly all configurations.

Taken together, these results indicate that deconvolution performance in BAL is determined as much by reference design and cell-type granularity as by the underlying algorithm. Linear and matrix factorization approaches generally achieve the highest accuracy when reference data are well matched to bulk samples, but their relative ranking can change substantially when the reference differs in chemistry, disease status, or preprocessing. Increasing resolution from broad lineages to macrophage-resolved sub-clusters amplifies the impact of reference mismatch and multicollinearity, leading many methods to fail at Level 3, whereas a small group of algorithms (CSCD/CSCD.F, Bisque/Bisque.F, CSsingle, AutoGeneS) retain moderate performance. The comparatively robust behavior of the in-study reference across all resolutions, and its slower performance decay in Fig. 4**D**, underscores the value of protocol-matched, pilot-scale single-cell profiling for accurate and stable deconvolution benchmarks.

### 2.4 A small in-study reference outperforms all external references across methods and resolutions

To quantify whether protocol-matched references improve deconvolution even when based on only a small number of samples, we compared the `in-study` reference—constructed from six randomly selected donors in each reverse cross-validation fold—against four external BAL scRNA-seq references using a unified linear mixed-effects model. For every method, annotation level, and replicate, we computed the paired difference

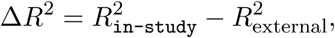

so that positive values directly quantify the advantage of the `in-study` reference. The estimated marginal means from this model are summarized in Fig. 5**A–C**.

Across all comparisons, the in-study reference produced consistently higher *R*^2^ values than every external reference (Fig. 5**A**). The advantage was large and statistically robust: estimated marginal means for the four external references ranged from 0.142 to 0.343, and all 95% confidence intervals excluded zero. These results demonstrate that even a small reference constructed from only six donors (20% of the cohort) provides substantially better deconvolution performance than larger public datasets that differ in chemistry, disease status, or preprocessing. Differences in marginal means across external datasets partly reflect method availability: some algorithms failed on particular references (e.g., DWLS on Liao and Li; CIBERSORTx on the BAL atlas), which depresses the average and highlights the challenges posed by heterogeneous public data.

The magnitude of the in-study advantage increased systematically with annotation resolution (Fig. 5**B**). At the broadest level (11 lineages), the average improvement was 0.148; this increased to 0.231 at intermediate resolution and 0.238 at the fine-grained Level 3. These substantial and highly significant fixed effects indicate that protocol matching becomes increasingly important as signature matrices become more collinear and cell-type distinctions more subtle. In other words, when the deconvolution task is hardest, a small but well-matched reference performs disproportionately better than large but heterogeneous external datasets. Performance gains varied across methods but remained positive for every algorithm tested (Fig. 5**C**). DWLS, HSPE, GEDIT, FARDEEP, and CSCD.F showed the strongest dependence on reference matching, with Δ*R*^2^ values exceeding 0.30, suggesting that these algorithms are particularly vulnerable to gene-level biases introduced by platform and processing differences. Other methodsincluding AutoGeneS (NNLS mode), CSsingle, and MuSiC.Fexhibited smaller but still positive differences (*<* 0.10), indicating greater robustness to reference heterogeneity. These method-specific patterns further emphasize that reference mismatch interacts with algorithmic design, amplifying platform-induced biases in some models more than others.

Taken together, the results in Fig. 5**A–C** demonstrate that even a six-donor `in-study` reference consistently outperforms all external references across methods, annotation levels, and replicates. Combined with our reverse 5-fold design, where every donor contributes exactly once to the reference and four times to evaluation, these findings show that only a modest investment in protocol-matched single-cell profiling is required to achieve substantial accuracy gains in real-bulk deconvolution benchmarks.

### 2.5 Paired samples enable discordant-gene detection and substantially improve deconvolution accuracy

In addition to providing a protocol-matched reference, even a small number of paired bulk single-cell samples offers a second advantage: they reveal genes whose expression is systematically inconsistent between true bulk RNA-seq and the pseudo-bulk profiles derived from the same donors. Such “discordant” genes violate the linear mixing assumption and disproportionately affect marker-driven deconvolution. Using the six paired donors available in each reverse cross-validation fold, we quantified bulk/pseudo-bulk log_2_ differences and applied four standard differential-expression frameworks (LIMMA, edgeR, DESeq2, paired *t*-test) to identify genes with reproducible discrepancies. We investigated whether the removal of these genes from both the reference matrix and bulk input enhances deconvolution performance across various methods, resolutions, and references (Fig. 6**A–C**).

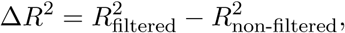

**Figure 5:**
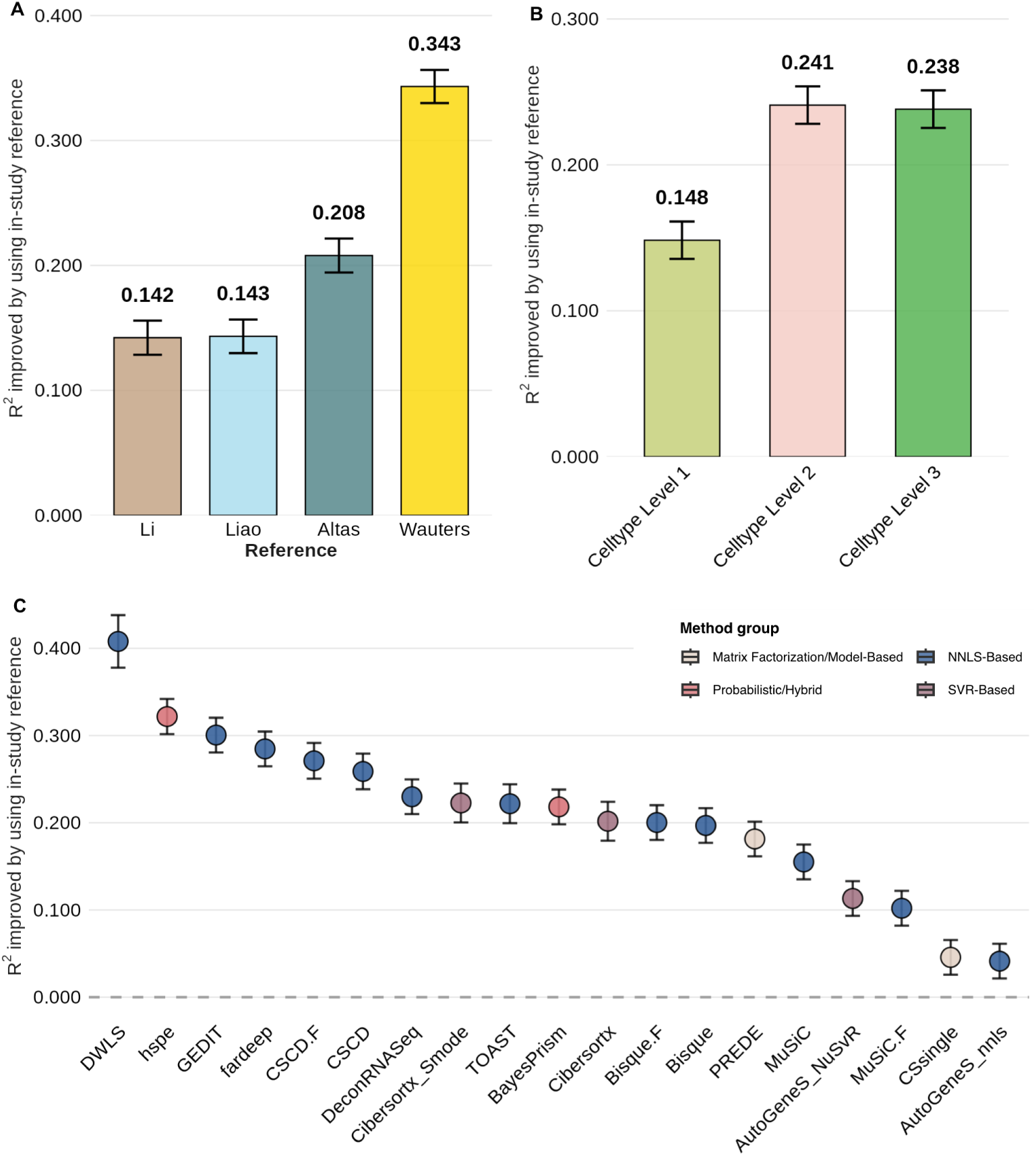
In-study vs. external reference performance comparison. (**A**) Reference dataset effects on *R*^2^ difference. The bar plot shows estimated marginal means (EMM) with 95% CI from LMER. Wauters *et al.*’s bar shows the largest gap (0.343), indicating the highest sensitivity to study-specific features. (**B**) Cell type resolution effects on *R*^2^ difference. EMM with 95% CI across three levels. Performance advantage increases with resolution: Level 1 (0.148), Level 2 (0.231), Level 3 (0.238), suggesting `in-study` references are more beneficial for fine-grained deconvolution. (**C**) Method-specific performance ranked by *R*^2^ difference. Horizontal bars show EMM with 95% CI for 15 methods. DWLS, hspe, GEDIT show the largest differences (>0.30), indicating strong dependence on in-study references. AutoGeneS nnls, CSsingle show the smallest differences (<0.10), suggesting better generalizability. Statistical model: LMER with fixed effects (method, level, reference) and random effects (patient, replicate, interaction). 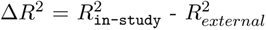. Positive values indicate `in-study` advantage.

**Figure 6:**
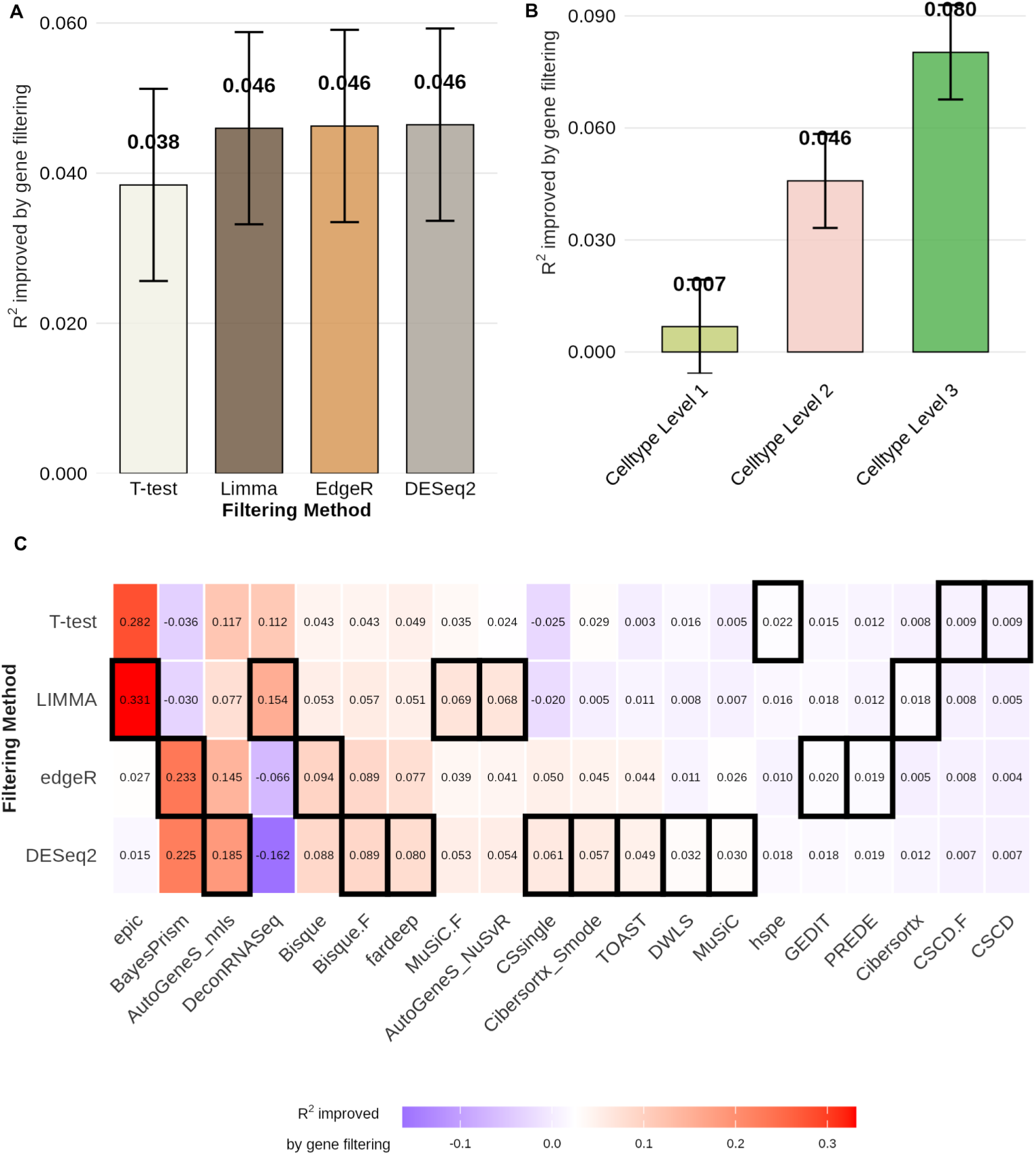
Removal of discordance genes between bulk and real-paired pseudo-bulk effects on deconvolution performance across methods and resolutions. (**A**) Filtering method effects on *R*^2^ difference. The bar plot shows estimated marginal means (EMM) with 95% CI from LMER. The LIMMA, edgeR, and DESeq2 show similar improvements (0.046), outperforming the paired *t*-test (0.038). (**B**) Cell type resolution effects on filtering benefit. EMM with 95% CI across three levels. Filtering benefit increases with resolution: Level 1 (0.007), Level 2 (0.046), Leve 3 (0.080), indicating fine-grained deconvolution benefits more from marker quality control. (**C**) Method-filter interaction effects. Heatmap showing the estimated marginal mean improvement in *R*^2^ (filtered - unfiltered) for each deconvolution method under each gene-filtering strategy. Cells indicate effect sizes, with warmer colors reflecting greater benefit from filtering. Black boxes highlight the best-performing filtering strategy for each method. Methods are ordered by their average filtering responsiveness. This interaction map demonstrates that no single filtering method is universally optimal; instead, sensitivity to filtering is strongly algorithm-dependent.

Filtering consistently improved accuracy across all statistical frameworks (Fig. 6**A**). Estimated marginal means from the mixed-effects model showed an average gain of ~ 0.04 in *R*^2^ after filtering, with LIMMA, edgeR, and DESeq2 performing similarly and outperforming the simple paired *t*-test. These improvements are modest in absolute magnitude but highly reproducible across the 636 evaluations, demonstrating that discordant-gene removal restores part of the model data alignment lost due to protocol differences between UMI-based single-cell libraries and bulk poly(A)^+^ sequencing.

The benefits of filtering increased sharply with annotation resolution (Fig. 6**B**). At the broad lineage level, gains were minimal, but at Levels 2 and 3 the average improvement rose to 0.046 and 0.080, respectively. This pattern reflects the rising signature collinearity and marker dependence at finer resolutions: when cell types are closely related, a handful of biased markers can dominate regression weights. Paired samples provide the only direct means to detect these genes, and their removal disproportionately stabilizes fine-grained deconvolution.

The magnitude and direction of improvement varied substantially across deconvolution methods and filtering strategies (Fig. 6**C**). No single differential-expression framework (LIMMA, edgeR, DESeq2, or paired *t*-test) consistently outperformed the others across all algorithms. Instead, each method exhibited a characteristic sensitivity profile: algorithms that rely heavily on free-form or partially constrained linear mixing, EPIC (Δ*R*^2^ = 0.33), AutoGeneS (NNLS mode) (Δ*R*^2^ = 0.18), and BayesPrism (Δ*R*^2^ = 0.23), showed the largest gains for their optimal filter, indicating a strong dependence on marker quality and a higher susceptibility to platform-induced gene-expression distortion. In contrast, methods with strong internal regularization or aggressive marker down-weighting (e.g. CSCD, DeconRNASeq, CIBERSORTx) displayed minimal changes (Δ*R*^2^ *<* 0.02), suggesting that their modeling assumptions buffer against discordant genes.

These heterogeneous responses demonstrate that discordant-gene filtering does not act as a generic preprocessing enhancement but interacts directly with the mathematical structure of each algorithm. Methods that treat all marker genes as equally informative benefit the most from removing genes that violate linear-mixing assumptions, whereas methods that internally shrink, reweight, or redefine gene sets gain relatively little. This method-specific landscape underscores the need for algorithm-tailored filtering strategies rather than a one-size-fits-all approach.

Taken together, these results show that paired samples serve a dual role in deconvolution studies. Beyond enabling a small but highly compatible in-study reference, they also provide the statistical power needed to identify and exclude discordant genes—an intervention that yields measurable and resolution-dependent gains across many algorithms. This dual utility underscores the value of generating even a modest number of paired bulk single-cell libraries in any deconvolution study where accurate fine-resolution inference is required.

Implement filtering as a standard preprocessing step, as it consistently enhances results, particularly when analyzing high-resolution cell types or employing methods that are known to yield benefits (EPIC, AutoGeneS nnls, BayesPrism). Check the method-specific standardization recommendations heatmap to confirm the interested deconvolution methods’ preference (refer to Fig. 6C).

### 2.6 Clinically relevant performance: recovering COPD-control differences

Previous sections demonstrated that (i) the `in-study` reference consistently outperforms external references, and (ii) discordant-gene filtering further improves accuracy. In this clinically oriented analysis, we therefore restrict attention to *only the in-study reference* and evaluate deconvolution methods *with filtering applied*. Our goal is to assess whether bulk deconvolution can reproduce the same COPD-control differences found using single-cell data and to determine which filtering strategies, cell-type resolutions, and deconvolution methods perform best in this setting.

For each cell type, we first used single-cell derived proportions to test association with COPD status (Wilcoxon rank-sum test, *p <* 0.05); these results define the true disease-associated cell types. We then applied deconvolution to the bulk RNA-seq data using the in-study reference and repeated the same association test. Treating the single-cell call (associated/not associated) as the true label and the bulk-based call as the prediction, we constructed a confusion matrix for every method, filtering strategy, and resolution, and computed precision (Fig. S3), recall (Fig. S4) and the *F*_2_ score (Fig. 7). The *F*_2_ metric up-weights recall and therefore reflects the requirement that clinically useful methods should avoid missing truly disease-associated cell types.

**Figure 7:**
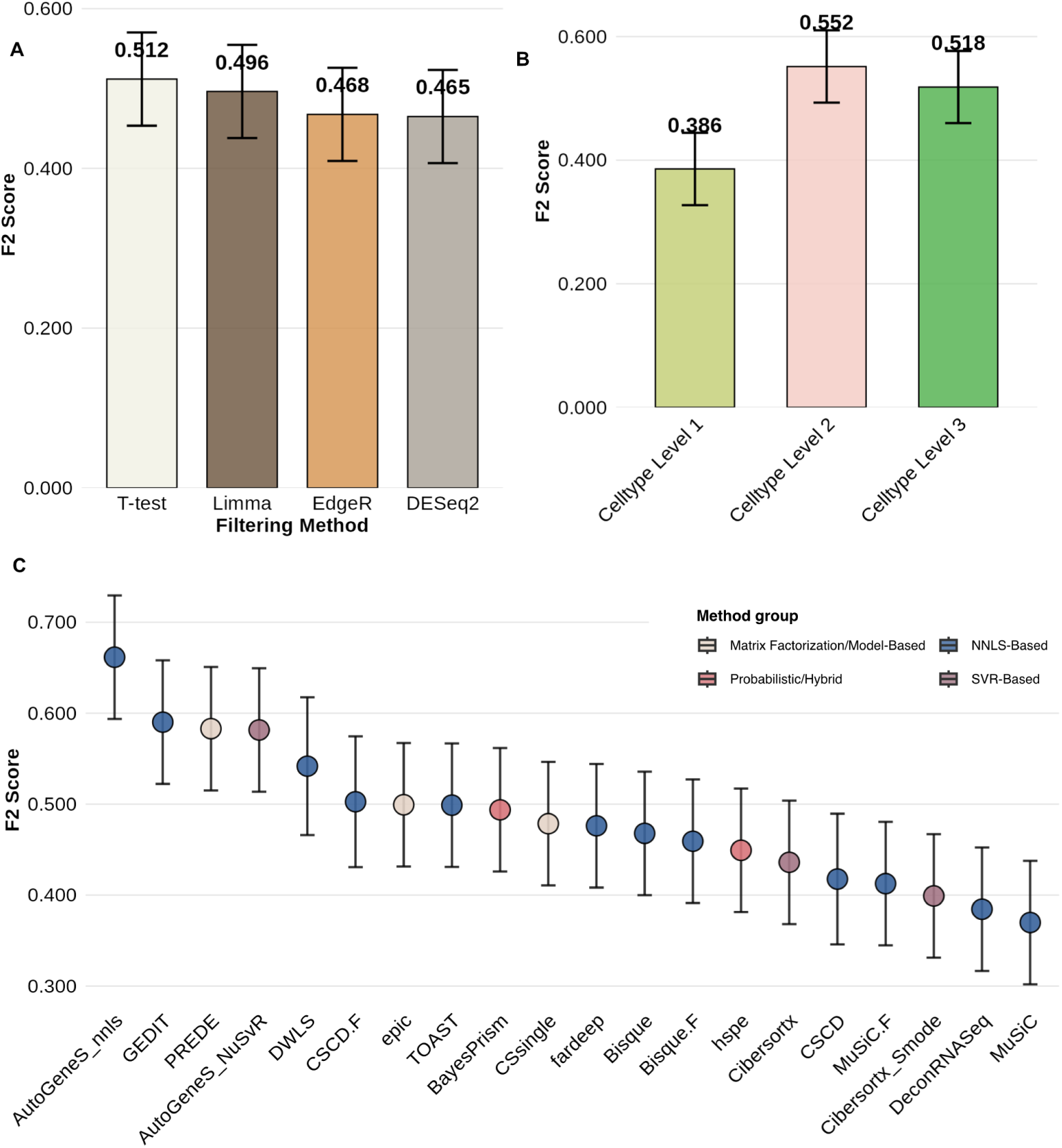
Clinically oriented evaluation using COPD status. (**A**) Effect of gene-filtering method on *F*_2_ score. (**B**) Effect of cell-type resolution on *F*_2_ score. (**C**) Comparison of deconvolution methods using the in-study reference under gene filtering.

#### Filtering strategy: sensitivity dominates

Across the four filtering strategies tested (paired *t*-test, LIMMA, edgeR, and DESeq2), gene filtering strongly influenced *F*_2_ and recall (Fig. 7**A**). Paired *t*-test filtering yielded the highest *F*_2_ (0.512) and recall (0.673), with LIMMA performing similarly. DESeq2 and edgeR produced significantly lower *F*_2_ scores (≈ 0.4650.468) and recall (≈ 0.5980.610). Marginal means were tightly clustered (0.290.30) across all filters, with no significant differences. Precision remained nearly unchanged across strategies (≈ 0.290.30), demonstrating that filtering strategies primarily modulate sensitivity (recall and F2), while positive predictive value remains stable across approaches. These results indicate that, when used with the `in-study` reference, simple variance-based filters (paired *t*-test or LIMMA) provide better clinical detection performance.

#### Cell-type resolution: moderate/fine granularity improves detection

We next evaluated which annotation resolution: broad (11 types), intermediate (13 types), or fine (18 sub-clusters), best supports identification of COPD-associated cell types using bulk data (Fig. 7**B**). *F*_2_ performance increased sharply from Level 1 (0.386) to Level 2 (0.552) and remained high at Level 3 (0.518). Precision showed a similar large improvement pattern between Level 1 and Levels 2-3. These trends indicate that moderate to fine-grained resolution provides more power for detecting disease-control contrasts than broad lineage categories. Among the three, Level 2 offered the strongest and most stable gains, balancing granularity with statistical robustness.

#### Deconvolution methods: Large variation in performance

Using the in-study reference and discordant-gene filtering, we compared fifteen deconvolution methods under each filtering strategy and resolution (Fig. 7**C**). Method choice was the dominant factor affecting clinical detection performance. AutoGeneS (NNLS) achieved the highest *F*_2_ (0.662) and led both recall and precision, followed by GEDIT, PREDE, and AutoGeneS (NuSVR mode). DWLS also performed well. In contrast, MuSiC and DeconRNASeq, despite strong compositional accuracy in earlier analyses, cannot show strong *F*_2_ score, recall, and precision in COPD association. This divergence highlights that methods optimized for stable proportion estimation are not necessarily optimal for detecting differential abundance associated with disease.

#### Summary

These findings complement the proportion-accuracy analyses and clarify which configurations are best suited for disease-focused studies. For studies aiming to detect disease-control differences in cell-type abundance, the optimal configuration is paired *t*-test or LIMMA filtering, moderate/fine cell-type resolution, and AutoGeneS (or GEDIT/PREDE as alternatives).

## 3 Discussion

To our knowledge, this study presents the first fully paired benchmark for transcriptomic deconvolution using human BAL samples, in which each donor contributes a protocol-matched bulk RNA-seq library and scRNA-seq profile. This design allows us to evaluate deconvolution performance on *real* bulk data, to disentangle the effects of algorithm choice and reference construction, and to examine how violations of the linear-mixing assumption influence both benchmarking outcomes and downstream biological inference.

Most existing benchmarks rely on pseudo-bulk aggregation of single-cell data, an approach that has enabled systematic method development and comparison at scale. Our results do not challenge the value of these studies. Rather, by leveraging paired data, we provide complementary evidence that clarifies when pseudo-bulk approximations are sufficient and when protocol mismatch, reference design, and gene-level discrepancies materially affect deconvolution results.

### Paired bulk–single-cell data reveal systematic but localized discrepancies

Across donors, real bulk RNA-seq and matched pseudo-bulk profiles show strong global concordance, confirming that pseudo-bulk aggregation is generally a high-fidelity approximation of bulk expression. At the same time, paired analysis reveals a reproducible subset of genes whose expression differs systematically between technologies. Although these genes represent a minority of the transcriptome, they include cell-type informative features commonly used in deconvolution signature matrices, such as canonical markers and genes with high leverage in regression-based models.

The direction and magnitude of these discrepancies are consistent with known protocol-specific effects, including poly(A)^+^ selection and fragmentation in bulk RNA-seq versus 3^′^-UMI capture in single-cell assays, gene-length sensitivity, and library construction differences. These factors introduce small but reproducible deviations from the linear-mixing assumption *b* = *Rw*. While such deviations have limited impact on global expression concordance, they disproportionately affect the genes that drive deconvolution, underscoring the unique value of paired data for quantifying technology-induced bias at functionally critical loci.

### Paired samples improve both reference construction and evaluation fidelity

Beyond characterizing bulk–pseudo-bulk concordance, paired data allow direct evaluation of how reference choice influences deconvolution inputs. By reconstructing reference-derived bulk profiles using multiple single-cell references and comparing them to observed bulk RNA-seq, we show that a protocol-matched `in-study` reference consistently yields bulk reconstructions that more closely resemble the true data than those derived from external datasets.

This improvement reflects two complementary effects. First, pairing improves construction of the signature matrix by aligning technical properties between *b* and *R*, particularly for high-loading marker genes. Second, it yields a cleaner evaluation setting in which discrepancies between predicted and observed bulk expression are less confounded by platform mismatch. These findings indicate that reference compatibility can outweigh reference size, especially for algorithms that rely directly on linear mixing.

### Reverse cross-validation demonstrates the power of small paired cohorts

A central methodological contribution of our study is the reverse five-fold cross-validation design, in which a small subset of paired donors is used to construct the reference, and the remaining donors are reserved exclusively for evaluation. This design provides two advantages. First, it ensures strict prevention of information leakage: no donors single-cell profile is ever used simultaneously as reference and ground truth. Second, it directly addresses a practical constraint faced by many studies-limited access to single-cell profiling.

Remarkably, we find that even six paired donors are sufficient to generate an in-study reference that outperforms substantially larger external references across methods and annotation resolutions. This result does not diminish the value of large public atlases; rather, it highlights how modest, targeted pairing within a study can deliver disproportionate gains by minimizing technical and biological mismatch.

### Paired data enable discordant-gene detection and algorithm-aware filtering

Paired samples offer a second, distinct advantage: they enable identification of genes whose expression is systematically discordant between true bulk and pseudo-bulk. Removing these genes from both the reference and bulk input improves deconvolution accuracy across most algorithms, with gains increasing at finer annotation resolutions where collinearity and marker overlap are greatest. Importantly, discordant-gene filtering interacts strongly with algorithmic design. Methods that rely heavily on unconstrained or weakly constrained linear mixing (e.g., EPIC, AutoGeneS in NNLS mode, BayesPrism) benefit most, whereas algorithms with strong internal regularization or marker down-weighting show limited sensitivity. No single differential-expression framework is universally optimal, emphasizing that filtering should be viewed as a method-aware optimization rather than a generic preprocessing step. To preserve fair comparisons, we apply discordant-gene filtering only when an in-study reference is available, maintaining an unfiltered baseline for external references.

### Clinical inference highlights distinct performance trade-offs

While proportion accuracy metrics such as MSE and *R*^2^ are informative, many deconvolution studies ultimately aim to identify cell types associated with disease or clinical outcomes. Using COPD status as a real clinical endpoint, we show that methods performing well in compositional accuracy do not necessarily excel at detecting disease-associated abundance changes. Algorithms that emphasize marker optimization and regression stability (AutoGeneS, GEDIT, PREDE, DWLS) achieve higher recall-weighted performance, whereas approaches optimized for robust composition estimation are less sensitive in this setting.

These results emphasize that the notion of a best method depends on analytic goals. Absolute compositional accuracy and sensitivity to group differences are related but distinct objectives, and benchmarking frameworks should reflect the intended downstream application.

### Implications, limitations, and future directions

Together, our findings show that paired bulk—single-cell designs complement pseudo-bulk based benchmarks by clarifying how reference compatibility, gene-level discrepancies, and evaluation strategy influence deconvolution performance. Rather than replacing existing approaches, paired benchmarks provide a principled way to calibrate them, identify method-specific sensitivities, and guide study-specific optimization.

Our study has several limitations. First, it focuses on BAL, an immune-rich compartment dominated by macrophages; extension to other tissues with different cellular architectures will be informative. Second, although reverse cross-validation provides stable estimates, the COPD subgroup is small, and the disease-association analysis should be viewed as illustrative. Third, we focused on RNA-seq and did not explicitly model absolute RNA content or non-linear compositional effects.

Future work could extend this framework to additional tissues, sequencing chemistries, and spatial transcriptomics, as well as explore algorithmic strategies that down-weight discordant genes implicitly rather than via explicit filtering. More broadly, our results suggest a practical and cost-conscious blueprint for improving deconvolution: generate a modest pilot of paired single-cell libraries, construct a protocol-matched reference, and leverage pairing to identify and mitigate technology-driven discrepancies. Even limited pairing can substantially improve both benchmarking rigor and biological interpretability.

## 4 Conclusions

Using split-sample, donor-matched bulk and scRNA-seq from human BAL, we establish a fully paired benchmarking framework for transcriptomic deconvolution and apply it to a comprehensive evaluation of fifteen methods across five single-cell references and three cell-type resolutions on *real* bulk RNA-seq data. Unlike benchmarks based solely on simulated mixtures, this design enables direct assessment of deconvolution performance under realistic experimental conditions, while a reverse cross-validation scheme ensures strict separation between reference construction and evaluation. Together, this framework provides a robust and leakage-free baseline for comparing algorithms, reference choices, and annotation granularity in immune-rich tissues.

Within this benchmark, we report three main empirical findings. First, real bulk and pseudo-bulk profiles exhibit systematic gene-level differences, including among cell-type informative genes that disproportionately influence deconvolution. Second, a protocol-matched in-study reference constructed from only a small fraction of donors consistently outperforms much larger external references, with the advantage becoming most pronounced at fine cell-type resolution. Third, paired samples enable the identification of reproducibly discordant genes whose selective removal further improves accuracy for many deconvolution methods. These effects are consistent across algorithms, resolutions, and evaluation metrics and are robust to alternative modeling and normalization choices.

Although motivated as a benchmark, the broader contribution of this work also providing practical guidance for study design. Our results show that generating single-cell data for a *small subset* of bulk samples-pilot pairing, can serve two critical purposes: constructing an in-study reference and diagnosing bulksingle-cell discordance for targeted gene filtering. With modest additional costs, this strategy delivers substantial gains in accuracy and stability, particularly in challenging high resolution settings, and reduces reliance on large, heterogeneous external references. We therefore propose pilot pairing as a general and cost-effective protocol for future bulk deconvolution studies. The real-paired dataset and open workflow presented here provide a foundation for extending these guidelines to other tissues, sequencing chemistries, and disease contexts.

## 5 Methods

### 5.1 Study design and sample processing

Thirty BAL specimens were obtained from the Centre for Heart Lung Innovation, Bronchoscopy Biobank, under an IRB protocol approved by the UBC/Providence Health Care Research Ethics Board (approval number: H15-01872). Immediately after collection, each BAL suspension was split 50/50: one half was pelleted for bulk RNA extraction, and the other half was kept on ice for single cell library preparation (Fig. 1). This paired aliquot design ensures that every bulk library has a protocolmatched single cell counterpart from the same donor, preserving real bulk biases while providing ground truth cell-type profiles.

#### Bulk RNA-seq

Total RNA was extracted with RNeasy Plus kit (QIAGEN, Stockache, Germany) (RIN > 8). The poly(A)^+^ libraries was prepared using standard protocol for the NEBnext Ultra ii Stranded mRNA (New England Biolabs, Ipswich, MA, USA). Libraries were sequenced on an Illumina NovaSeq 6000 with samples having an average depth of 23.2 millions reads (42 bp x 42 bp reads). Reads were trimmed and quality filtered with FASTX Toolkit, aligned to Human reference with version GRCh38.p14 via STAR (v2.7.11a) [24], and quantified at the gene level with RSEM (v1.3.3) [25] using GENCODE v46 gene annotations. Bulk RNA-seq data contained 63,187 genes across 30 samples. For deconvolution benchmarking, 11,397 genes common to both bulk and paired single-cell datasets were used.

#### Single-cell RNA-seq

Single-cell libraries were generated with 10x Genomics Chromium Single-Cell 3’ v3.1 chemistry, targeting ~10,000 cells per aliquot. Sequencing was performed on the same Illumina platform using 2 × 150 bp paired-end reads, with a median depth of 40,000 reads per cell. Raw FASTQs files were processed using CellRanger (v8.0.0) with the human reference genome GRCh38.p14 and GENCODE v46 gene annotations to generate genecell count matrices. Cells expressing *<* 200 genes, *>* 2,500 genes, or *>* 5% mitochondrial counts were excluded. Using Seurat v5, UMI counts were log-normalized, 2,000 highly variable genes were selected, and standard principal component analysis (PCA) was used for dimensionality reduction. After retaining only genes common across all samples, 13,915 genes remained, with per-donor recoveries of 346-29,076 cells (median 5,965). The processed bulk RNA-seq and single-cell RNA-seq data - referred as “in-study” dataset - are available in the Gene Expression Omnibus (GEO) under accession number XXX.

### 5.2 Reference datasets and preprocessing

Robust benchmarking requires testing how deconvolution behaves when the single-cell reference differs in biology, chemistry, or processing from the bulk cohort. We therefore analyzed five references that span a compatibility gradient: three external BAL scRNA-seq studies, a harmonized multi-study BAL atlas, and an in-study reference created from a random 20% subset of 30 aliquots derived from our split-sample donors. Analyses were performed using a unified gene set intersected across all datasets. Quality filtering steps have been applied to all datasets that lacked a processed version. Each deconvolution method was conducted in accordance with its official tutorials and input requirements. For the five scRNA-seq references, uniform normalization and transformation to counts per million (CPM) were applied to mitigate batch effect and technical variability, following the guidelines of each deconvolution method. Reference and bulk datasets were then organized into the appropriate format, ensuring consistency in workflow and parameter settings.

#### External BAL scRNA-seq datasets

##### Li *et al.* 2022 Cystic-FibrosisHealthy BALF (GSE193782)

Seven donors (4 healthy, and 3 with cystic fibrosis) were profiled with 10x Genomics v3 chemistry, generating data from 113,213 cells and yielding 14 major immune and epithelial populations [26]. We downloaded the processed count matrix and metadata from GEO for further analysis.

##### Liao *et al.* 2020 COVID-19 severity cohort (GSE145926)

This dataset contains nine COVID-19 patients (6 severe, and 3 moderate) and three healthy controls. The single-cell libraries of these samples were prepared using 10x v2 chemistry[27] and the processed single-cell dataset, including [~63 103] cells after quality control filtering, were downloaded from UCSC CellBrowser (http://cells.ucsc.edu/covid19-balf/nCoV.rds).

##### Wauters *et al.* 2021 Mild/critical COVID-19 and non-COVID pneumonia

The single-cell libraries from 37 donors (24 patients with COVID-19, and 13 with pneumonia) were profiled using the 10x Genomics v3 chemistry. After quality-control filtering, a total of 120,871 high-quality cells were retained, and 20 immune and epithelial subclasses were annotated[28]. The processed counts matrix dataset was downloaded from the study [28] for further analysis.

##### Lung BAL scRNA-seq atlas (BAL-EA)

We pooled healthy-control subsets from several published BAL studies and reprocessed them through a unified Seurat v3 pipeline, including ambient-RNA removal, selection of 3 000 highly variable genes (HVGs), and reciprocal PCA integration). Across n = 26 healthy donors, this yielded a dataset of 352,732 cells.

#### In-study real-paired BAL scRNA-seq dataset

We processed in-study BAL samples from 32 donors (4 COPD and 28 non-COPD). After quality control, two non-COPD scRNA-seq samples were removed, yielding 241,924 high-quality single cells. Because these libraries share collection, preservation, and sequencing protocols with the bulk cohort, they minimize batch effects and technology-specific biases.

##### Uniform preprocessing pipeline

All five datasets were processed through one Seurat v3 workflow: select overlapping genes among all references, CPM and log-normalization based on the input requirement of deconvolution methods, and selection of highly variable genes for Cibersortx. Gene symbols were taken from the intersection of all scRNA-seq datasets, and matrices were trimmed to the bulk-single-cell intersect of 11,397 genes.

### 5.3 Cell-type annotation and resolution hierarchy

Cell-type labels were harmonized across our in-study paired data and all references using the reannotation labels from Hu *et al.* [23], which were designed to ensure cross-dataset comparability in BAL scRNA-seq. For each dataset, we mapped author-provided labels to this unified taxonomy and manually resolved naming mismatches (e.g., macrophage/monocyte states) to preserve consistent biological interpretation across studies.

Every reference was re-annotated with a shared three-level hierarchy to quantify how granularity influences accuracy: **(1.) 11 broad cell types** B cell, CD4^+^ Tcell, CD8^+^ Tcell, conventional dendritic cell, epithelial cell, macrophage, neutrophil, mast cell, NK cell, plasmacytoid dendritic cell, and plasma cell. **(2.) 13 intermediate types** subdivision of macrophage (alveolar macrophage, non-alveolar macrophage, monocyte), yielding 13 categories. **(3) 18 fine sub-clusters** further splitting of non-alveolar macrophage states (CCL18^+^, CCL2^+^, RGS1^+^, MT1G^+^), monocyte subsets (FCN1^+^, IL1B^+^).

### 5.4 Signature matrix generation and marker genes

For every reference-resolution combination, we use the feature gene list from Hu *et.al* [23], which first manually maps common cell types from the HLCA core and Wauters’ BAL dataset [28] for cross-dataset comparability. To prevent batch-related artifacts, DE analyses were performed individually in each HLCA v1.0 core sub-study (n = 11) and resampled control subsets of the Wauters BAL dataset. Seurat’s FindAllMarkers tool (“MAST”) found cell-type-specific DE genes for each research, retaining genes with an adjusted *p*_*value <* 0.05 and |log_2_ *FC*|≥ 1. DE genes were rated by repeatability across datasets for each cell type, and genes found in multiple independent analyses were retained at a threshold. From genes recurring in ≥10 datasets, thresholds were calculated hierarchically and relaxed sequentially (≥9, ≥8, …, ≥3) until each cell type produced an acceptable marker panel (*>*5 genes). Upon eliminating duplicate genes across various cell types, the marker list included 664 genes at Level 1, 723 at Level 2, and 851 at Level 3.

### 5.5 Deconvolution methods evaluated and execution rules

To ensure a fair and reproducible comparison, all deconvolution methods were executed under explicitly defined and method-consistent input, preprocessing, and parameterization rules. This subsection defines the algorithmic scope of the benchmark and standardizes how each method interfaces with bulk and single-cell data, independent of reference choice, cross-validation design, or gene filtering.

#### 5.5.1 Methods to be evaluated

To provide an up-to-date performance landscape, we benchmarked **fifteen** representative algorithms released between 2013 and 2025 (Table 2) in recent peer-reviewed benchmarking studies.

**Table 2:** Overview of the fifteen deconvolution algorithms benchmarked.

| Method | Core algorithm | Supervision | Input genes* | ScRNA ref. | Primary application |
| --- | --- | --- | --- | --- | --- |
| *NNLS / linear models* |  |  |  |  |  |
| Bisque | NNLS + gene-wise transform | Sup. | both | yes | Adipose, Cortex tissue |
| CSCD | shrinkage-covariance NNLS | Semi | both | yes | Embryonic stem, cortex tissue |
| DeconRNASeq | quadratic NNLS | Sup. | both | no | Mixed |
| DWLS | dampened WLS | Sup. | auto | yes | Blood, Mouse mixed |
| GEDIT | NNLS | Semi | auto | yes | Mixed, Ascites, Blood |
| FARDEEP | aLTS-filtered NNLS | Sup. | both | no | Tumour |
| MuSiC | wNNLS | Sup. | both | yes | Pancreatic islets |
| MuSiC2 | iterative wNNLS | Sup. | auto | yes | Disease-mismatch |
| TOAST | OLS + iter. feature sel. | Unsup. | auto | yes | Rheumatoid arthritis, Breast |
| *SVR* |  |  |  |  |  |
| CIBERSORTx | ν-SVR | Sup. | auto | yes | Tumour, Blood |
| AutoGeneS | MOO-optimised SVR | Sup. | auto | yes | Embryonic, Blood |
| *Probabilistic / hybrid* |  |  |  |  |  |
| BayesPrism | Dirichlet multinomial | Unsup. | auto | yes | Tumour |
| HSPE | hybrid linearlog GLM | Sup. | auto | yes | blood, mixed |
| *Matrix / model-based* |  |  |  |  |  |
| EPIC | constrained LS + mRNA scaling | Sup. | both | no | Tumour, Blood |
| PREDE | partial-ref NMF | Semi | both | yes | Tumour, Blood |
| CSsingle | robust NMF (bulk mode) | Semi | auto | yes | Gastroesophageal tissue |

\*Input genes indicates whether the method consumes a user-supplied signature list (list), selects markers internally (auto), or both. \* GEDIT and AutoGeneS based on python, CIBERSORTx provide a web-tool, others are R packages.

##### Implementation and run rules

For every tool, we used the original authors implementation (R or Python); default hyperparameters were retained unless the documentation mandated changes for scRNA-seq references (Github: https://github.com/yushannn/BALbenchmark.git). All methods that accept a signature gene list were executed under two feature selection protocols that mirror common practice: **(1.) Signature gene selected**, for details, see Section 5.4; **(2.) Automatic detected gene**, use the default configuration of methods, letting it automatically select/weight genes.

##### Family I — Non-negative least squares (NNLS)

Classic linear mixing is implemented by DeconRNASeq [12], while MuSiC [13] and Bisque [29] weight genes to account for donor-to-donor heterogeneity or single-cell/bulk technology bias. CSCD extends NNLS with covariance regularization to dampen collinearity [30]. DWLS introduces adaptive gene weights for rare types [31], whereas FARDEEP employs an aLTS outlier filter well suited to tumor samples [32]. GEDIT uses entropy-based signature gene selection and non-negative linear regression [33].

##### Family II — Support vector regression (SVR)

CIBERSORTx couples *ν*-SVR with single-cell-guided batch correction and permits imputation of cell-type-specific expression profiles [2]. AutoGeneS replaces manual marker curation by optimizing the SVR objective over gene-set space [15].

##### Family III — Probabilistic and hybrid models

BayesPrism performs Bayesian deconvolution under a dirichlet-multinomial hierarchy and jointly refines reference expression [14]. MuSiC2 iteratively removes condition-specific DE genes to adapt references across disease states [34]. HSPE fits a hybrid linear/log-scale model that captures mean-variance coupling in RNA-seq data [35]. HSPE is the updated version of dstangle [36].

##### Family IV — Matrix-factorisation / model-based

TOAST (unsupervised) alternates between proportion estimation and feature selection [37]; PREDE integrates partial reference profiles into non-negative matrix factorization [38], and EPIC models cell-type-specific mRNA content to accommodate uncharacterized other cells in tumors [39]. CSSingle generalizes non-negative factorization to the spatial-omics setting but can be run in the bulk model [40].

#### 5.5.2 Input requirements and preprocessing to satisfy method interfaces

These methods differ substantially in their input data requirements, preprocessing strategies, and algorithmic approaches (Table 3).

**Table 3:**
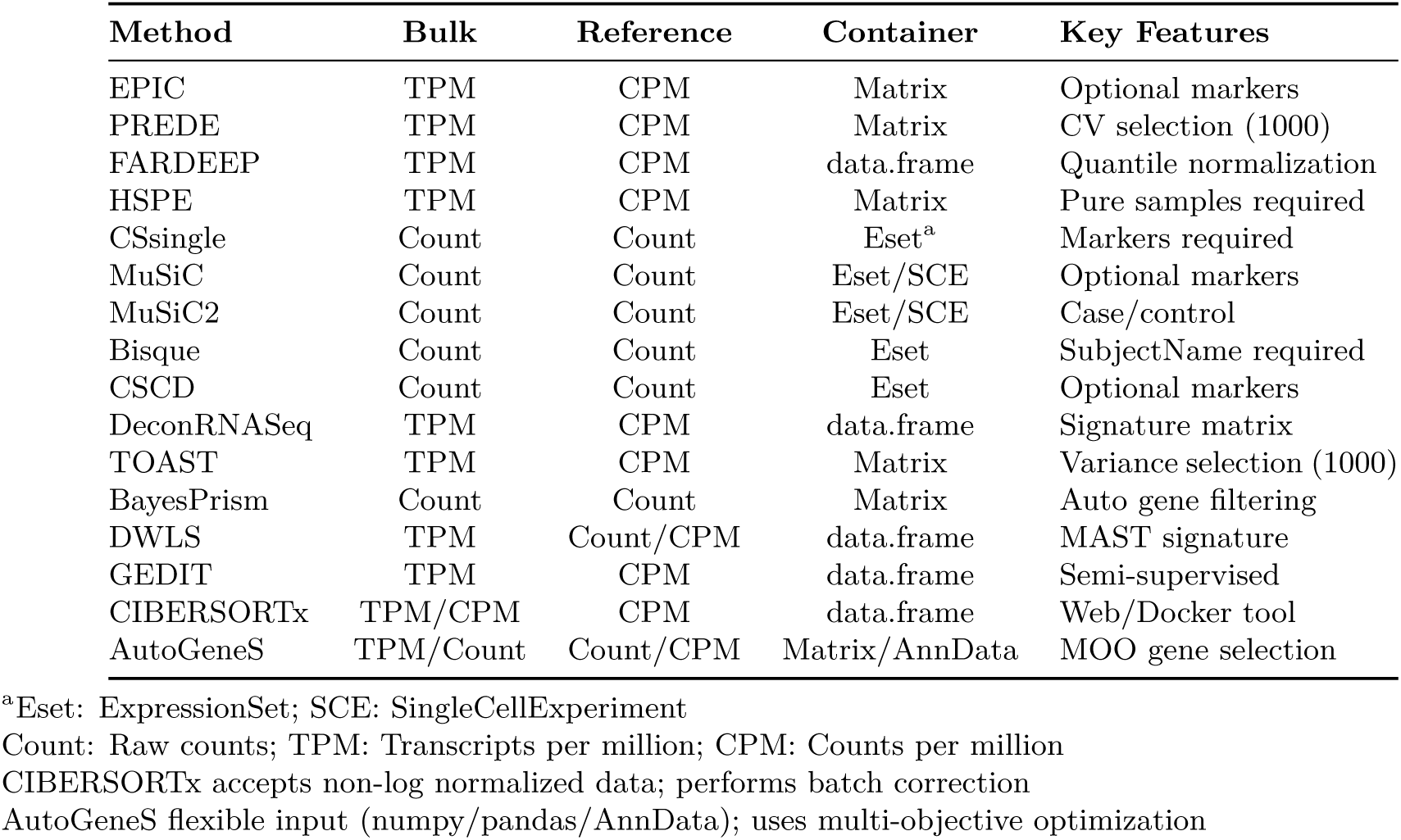
Summary of deconvolution methods and input requirements.

| Method | Bulk | Reference | Container | Key Features |
| --- | --- | --- | --- | --- |
| EPIC | TPM | CPM | Matrix | Optional markers |
| PREDE | TPM | CPM | Matrix | CV selection (1000) |
| FARDEEP | TPM | CPM | data.frame | Quantile normalization |
| HSPE | TPM | CPM | Matrix | Pure samples required |
| CSsingle | Count | Count | Eset<sup>a</sup> | Markers required |
| MuSiC | Count | Count | Eset/SCE | Optional markers |
| MuSiC2 | Count | Count | Eset/SCE | Case/control |
| Bisque | Count | Count | Eset | SubjectName required |
| CSCD | Count | Count | Eset | Optional markers |
| DeconRNASeq | TPM | CPM | data.frame | Signature matrix |
| TOAST | TPM | CPM | Matrix | Variance selection (1000) |
| BayesPrism | Count | Count | Matrix | Auto gene filtering |
| DWLS | TPM | Count/CPM | data.frame | MAST signature |
| GEDIT | TPM | CPM | data.frame | Semi-supervised |
| CIBERSORTx | TPM/CPM | CPM | data.frame | Web/Docker tool |
| AutoGeneS | TPM/Count | Count/CPM | Matrix/AnnData | MOO gene selection |

<sup>a</sup>Eset: ExpressionSet; SCE: SingleCellExperiment
Count: Raw counts; TPM: Transcripts per million; CPM: Counts per million
CIBERSORTx accepts non-log normalized data; performs batch correction
AutoGeneS flexible input (numpy/pandas/AnnData); uses multi-objective optimization

##### Bulk RNA-seq data normalization

The methods can be categorized into two groups based on bulk data input requirements. Raw count-based methods (MuSiC, MuSiC2, Bisque, CSCD, CSsingle, and BayesPrism) require unnormalized read counts and implement internal normalization procedures to account for library size and technical variations. TPM-normalized methods (EPIC, PREDE, FARDEEP, HSPE, DeconRNASeq, TOAST, and DWLS) require transcripts per million normalized expression values, which account for both sequencing depth and gene length.

##### Reference single-cell data preparation

Reference single-cell RNA-seq data preprocessing also varies by method. Methods using counts per million normalization (EPIC, PREDE, FARDEEP, HSPE, DeconRNASeq, and TOAST) require CPM-normalized reference profiles, calculated by dividing raw counts by total reads per cell and multiplying by one million. Methods using raw counts (MuSiC, MuSiC2, Bisque, CSCD, CSsingle, and BayesPrism) utilize unnormalized count matrices directly.

##### Data container and format requirements

Methods require different data structures for computation. Several methods (CSsingle, MuSiC, Bisque, CSCD, and DeconRNASeq) require Bioconductor ExpressionSet objects [41]. MuSiC and MuSiC2 additionally require SingleCellExperiment [42] objects for reference data. The remaining methods (EPIC, PREDE, FARDEEP, HSPE, TOAST, DWLS, and BayesPrism) accept standard R matrices or data frames. Specific metadata requirements vary: MuSiC, MuSiC2, Bisque, and CSCD require subject identifiers; Bisque and CSCD additionally require explicit cell type labels in the metadata.

### 5.6 Reverse five-fold cross-validation benchmark framework

#### Design rationale

The design of our benchmarking framework is motivated by two complementary objectives. First, to provide a statistically reliable and realistic benchmark for transcriptomic deconvolution, we generated a large paired dataset comprising 30 donors and over 240,000 single cellsa scale that is substantial by contemporary standards for human single-cell studies, even without paired bulk RNA-seq. This scale enables robust estimation of method performance on real bulk data while capturing biological and technical heterogeneity across donors.

Second, beyond benchmarking, we seek to address a practical question faced by many bulk transcriptomic studies: whether accurate deconvolution requires a large single-cell cohort or whether a small number of paired samples can already provide a high-quality reference. These two objectives (maximizing benchmarking robustness while evaluating data efficiency) jointly motivate our use of a reverse cross-validation strategy.

#### Design of the computer experiment

To reconcile these goals, we implemented a donor-level reverse five-fold cross-validation scheme (Fig. 1). The design is termed reverse because, unlike conventional benchmarking approaches that fix a large single-cell reference and cross-validate only bulk samples, we repeatedly reconstruct the in-study single-cell reference from a small subset of paired donors while evaluating performance on the remaining bulk samples.

Specifically, the 30 paired BAL donors were randomly partitioned into five folds of six donors each. In each iteration, non-COPD donors in the held-out fold were used exclusively to construct the `in-study` scRNA-seq reference and corresponding signature matrix *R*. The remaining 24 donors served as evaluation samples: their bulk RNA-seq profiles were deconvolved to obtain estimated cell-type proportions *ŵ*.

For evaluation, ground-truth cell-type proportions *w* were computed independently for each evaluation donor and annotation level by counting cells assigned to each cell type in the donors paired scRNA-seq data and normalizing by the total number of high-quality cells. These ground-truth proportions were used solely for benchmarking and never contributed to reference construction within the same fold. Because reference donors and evaluation donors are disjoint in every iteration, no donors single-cell data is used simultaneously for reference construction and performance evaluation, ensuring leakage-free assessment.

By iterating over the five folds, each donor contributes once to reference construction and four times to evaluation, yielding five independent replicates per method. Importantly, this design allows us to quantify how well a reference constructed from only six paired donors (20% of the cohort) performs relative to substantially larger external references, while still leveraging the full paired dataset to obtain stable and reproducible performance estimates.

External single-cell references were evaluated on identical bulk splits to isolate reference effects from partition variability.

Within this reverse cross-validation framework, we additionally assessed the impact of discordant-gene filtering when an in-study reference was available. Specifically, for each fold, all deconvolution analyses using the in-study reference were performed both with and without gene filtering, allowing the effect of filtering to be quantified independently of reference construction.

Across five folds and three cell-type resolutions, we benchmarked 15 deconvolution methods (14 implemented in `R` and CIBERSORTx accessed via its web interface) under baseline and filtered conditions. In total, this design yielded

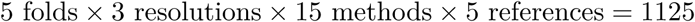

baseline runs, and

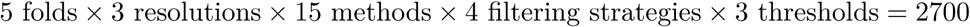

filtered runs, for a total of 3825 deconvolution evaluations.

Discordant-gene filtering was applied exclusively to the in-study reference, because gene-level discordance can only be quantified when matched bulk-single-cell pairs are available. External references were evaluated on the same bulk splits without filtering to isolate reference-dependent effects. The definition and implementation of discordant-gene filtering are described in detail in the following subsection.

### 5.7 Discordant-gene detection and filtering

Most transcriptomic deconvolution methods model bulk RNA-seq expression as a linear mixture of cell-typespecific reference profiles. In practice, this assumption can be violated for a subset of genes whose expression is systematically distorted by protocol differences between bulk and single-cell RNA-seq, including polyA^+^ selection versus 3^′^-end UMI capture, gene-length effects, RNA degradation, and alignment or quantification biases. When such genes carry high weight in reference signatures, they can disproportionately affect deconvolution accuracy.

Leveraging the availability of protocol-matched bulk and single-cell data, we implemented a paired-sample discordant-gene filtering strategy that explicitly identifies and removes genes exhibiting reproducible cross-modality discordance. This procedure was applied exclusively to in-study references, where direct donor-level comparison between bulk and pseudo-bulk is possible. To isolate its contribution, all analyses using the in-study reference were performed both with and without filtering.

#### Pseudo-bulk construction

Within each cross-validation fold, pseudo-bulk profiles were generated from the six donor scRNA-seq samples used to construct the in-study reference. For each donor, single-cell UMI counts were depth-normalized to CPM and aggregated by averaging across cells. Bulk RNA-seq expression was length-normalized to TPM. Both bulk and pseudo-bulk profiles were transformed as log_2_(*x* + 1), yielding log_2_(TPM + 1) for bulk and log_2_(CPM + 1) for pseudo-bulk. This procedure aligns the gene universe while mitigating gene-length bias present only in bulk RNA-seq.

#### Detection of discordant genes

Within each fold, donor-matched bulk and pseudo-bulk profiles were compared using two complementary analysis branches. (1.) Normalized-expression branch: bulk and pseudo-bulk data were standardized using either Z-score normalization alone or log_2_(*x*+1) transformation followed by Z-scoring. Paired *t*-tests and LIMMA were applied to detect systematic differences, with thresholds |Δ*Z*| *>* 0.5, 1.0, 1.5 (Z-score only) or |Δ*Z*| *>* 1.0, 1.5, 2.0 (log_2_+Z). (2.) Raw-count branch: raw UMI counts were summed per donor to construct pseudo-bulk count matrices. Differential expression was assessed using edgeR (TMM normalization, quasi-likelihood framework) and DESeq2 (size-factor normalization with shrinkage), with thresholds |log_2_ FC| *>* 1.5, 2.0, 3.0.

All tests used Benjamini-Hochberg correction with FDR control at 0.05, resulting in 12 filtering strategies (4 statistical methods × 3 thresholds).

#### Filter application

Within each fold, genes flagged as discordant by any filtering strategy were removed from both the reference/signature matrix and the corresponding bulk expression profiles prior to deconvolution. This ensures that proportion estimates are derived from genes that are consistent across bulk and single-cell modalities within the same experimental context.

#### Scope and rationale

Discordant-gene filtering was restricted to in-study references, where protocol-matched bulk-single-cell pairs permit direct quantification of technical discordance. External references and the BAL atlas were left unfiltered, both because such pairing is unavailable and to provide a neutral baseline for evaluating reference-dependent effects. This design allows the added value of paired data, beyond reference construction alone, to be isolated and quantified.

### 5.8 Evaluation metrics and statistical models

Let *w_ij_* denote the *true* proportion of cell type *j* ∈ {1, …, *K*} in bulk sample *i* ∈ {1, …, *n*}, derived from paired single-cell RNA-seq, and let *ŵ_ij_* denote the corresponding estimate returned by a deconvolution method.

#### 5.8.1 Performance metrics for proportion estimation

We quantified deconvolution accuracy using complementary metrics that capture absolute error, explained variance, and rank concordance.

**Mean squared error (MSE).**

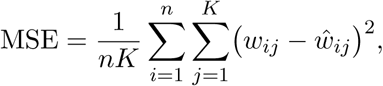

where *n* = 24 evaluation samples per cross-validation fold and *K* is the number of annotated cell types (11, 13, or 18). MSE emphasizes absolute deviation and is sensitive to large errors in rare cell types.

**Coefficient of determination (*R*^2^).**

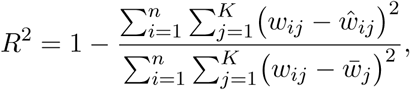

where *w̄_j_* denotes the mean proportion of cell type *j* across donors. *R*^2^ captures explained variance and enables direct comparison across methods, references, and annotation resolutions.

##### Correlation metrics

Pearson and Spearman correlations between *w* and *ŵ* were computed as secondary diagnostics to assess linear and rank concordance, respectively. These metrics are reported in the Supplementary Material and used for sensitivity analyses.

#### 5.8.2 Hierarchical statistical modeling

To quantify systematic effects while accounting for repeated measures across donors, folds, methods, and references, we employed linear mixed-effects models (LMER) fit using restricted maximum likelihood (REML) via the lme4 and lmerTest packages.

##### Model 1: Reference dataset effects

To assess the advantage of protocol-matched references, we modeled paired differences in explained variance:

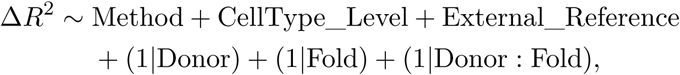

where 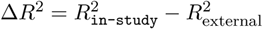

##### Model 2: Discordant-gene filtering effects

To evaluate filtering strategies, we modeled performance differences between filtered and unfiltered analyses for the in-study reference:

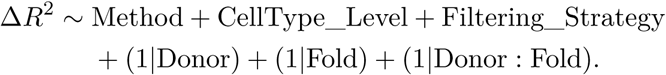

##### Model 3: Clinically oriented performance

For disease-association analyses, we modeled the *F*_2_ score:

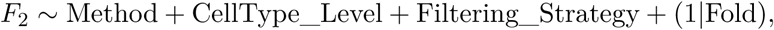

prioritizing recall to reflect the cost of failing to detect disease-associated cell types.

#### 5.8.3 Statistical inference and uncertainty quantification

Estimated marginal means (EMMs) and 95% confidence intervals were obtained using the `emmeans` package. Pairwise contrasts between references or filtering strategies were evaluated using two-sided paired *t*-tests and Wilcoxon signed-rank tests, with *α* = 0.05. Wilcoxon tests were used for primary inference due to robustness against non-normality.

#### 5.8.4 Classification metrics for clinical evaluation

For cell-typelevel disease association, we constructed confusion matrices and computed:

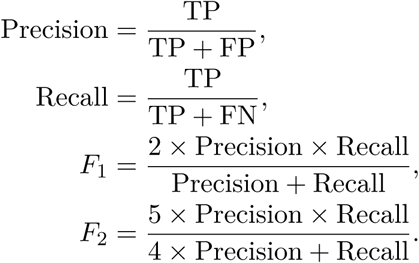

The *F*_2_ score upweights recall (*β* = 2), emphasizing sensitivity in detecting clinically relevant cell-type differences.

### 5.9 Computational environment

All analyses were performed on Compute Canada’s Digital Research Alliance high-performance computing infrastructure using StdEnv/2020. Core statistical analyses and deconvolution bench-marking were conducted in R (version 4.3.1), employing the following key packages: Seurat (v 4.3.0) for single-cell data processing; limma (v3.58.1), edgeR (v4.0.16), and DESeq2 (v1.42.1) for differential expression and discordant-gene filtering; lme4 (v1.1.35.5) and lmerTest (v3.1.3) for mixed-effects modelling; and tidyverse (v2.0.0) for data manipulation. Fifteen deconvolution methods were benchmarked, including EPIC (v1.1.7), FARDEEP (v1.0.1), MuSiC (v1.0.0), MuSiC2 (v0.1.0), BisqueRNA (v1.0.5), BayesPrism (v2.2.2), DWLS, hspe (v0.1), TOAST (v1.16.0), CSC-DRNA (v1.0.3), CSsingle (v0.1.0), DeconRNASeq (v1.44.0), and PREDE (v1.2.1). CIBERSORTx deconvolution was performed via its web interface (https://cibersortx.stanford.edu/). Additional methods, AutoGeneS and GEDIT, were implemented in Python (version 3.10) with numpy, pandas, and scipy.

Visualization was performed using ggplot2 (v3.5.1), patchwork (v1.2.0), and cowplot (v1.1.3). To ensure reproducibility, all random seeds were fixed.

## Authors contributions

Conceptualization, X.Z. and D.D.S.; funding acquisition, X.Z., D.D.S., and X.S.; supervision, X.Z., and X.S; methodology, Y.H., X.S. and X.Z.; investigation, Y.H.; data curation, Y.H., Z.L. and D.T.; formal analysis, Y.H. and D.T.; writing original draft, Y.H.; writing review & editing, all authors

## Acknowledgments

We thank the Computational resources that were provided by the Digital Research Alliance of Canada (the Alliance). Fig. 1 was created with BioRender.com.

## Ethics approval and consent to participate

UBC/Providence Health Care Research Ethics Board (approval number: H15-01872). For data links, please upload de-identified data onto GEO.

## Supplementary Information

**Figure S1:**
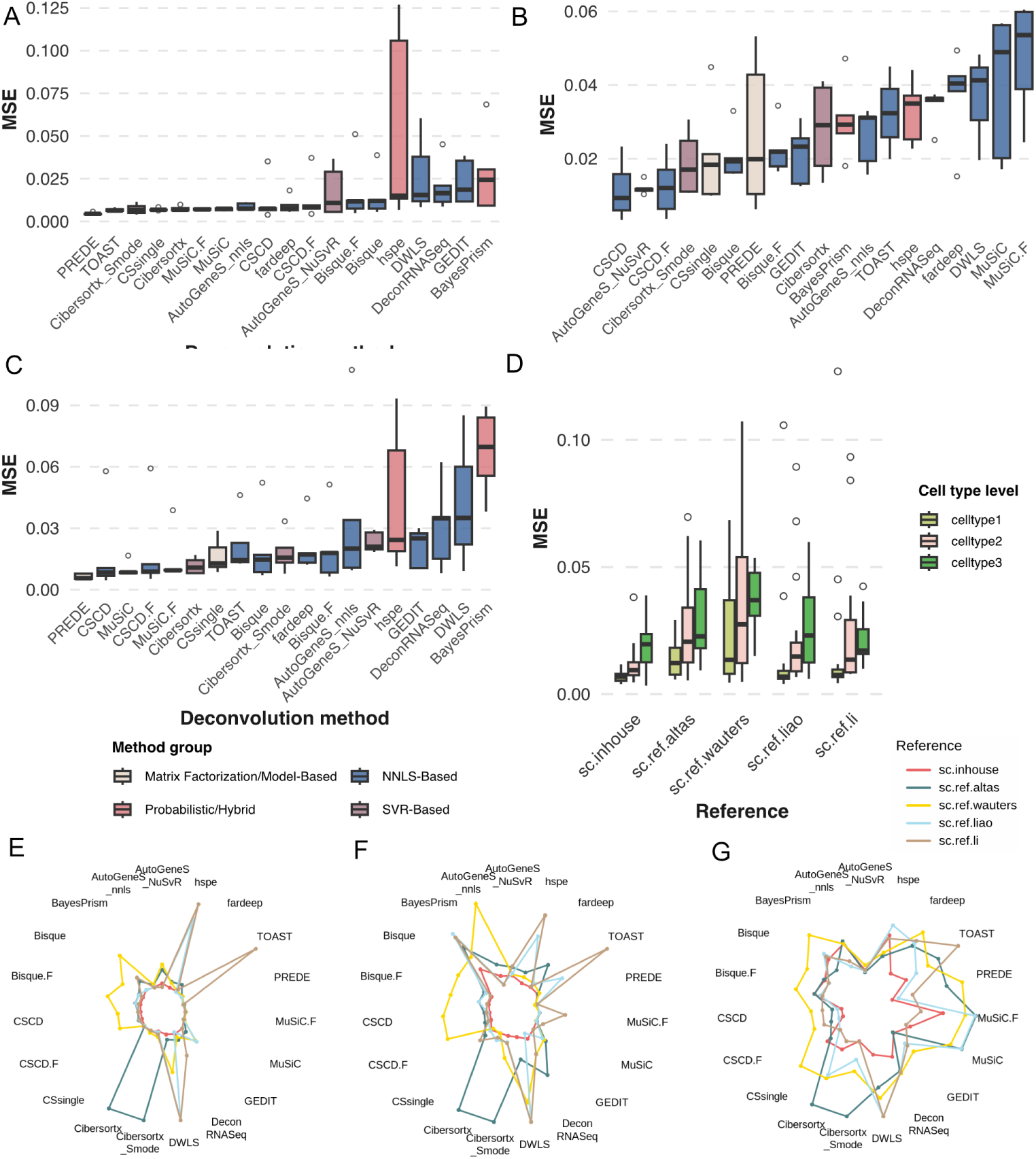
Comprehensive evaluation of deconvolution method performance across three levels and five references. (**A—C**) *MSE* performance by the deconvolution method from Level 1 to Level 3. Boxplots show distribution across reference datasets, colored by method group. Methods ordered by median performance. (**D**) Cross-resolution performance comparison. Boxplots show *MSE* distributions across alternative references (gray/beige/green for Levels 1/2/3). (**E—G**) Radar charts comparing method performance across five reference datasets at Level 1, 2, and 3, respectively. Colored lines represent references: `in-study` (pink), BAL atlas (teal), Liao *et al.* (blue), Wauters *et al.* (yellow), and Li *et al.* (brown). Radial distance indicates *MSE* value. Polygon size increases with finer resolution.

**Figure S2:**
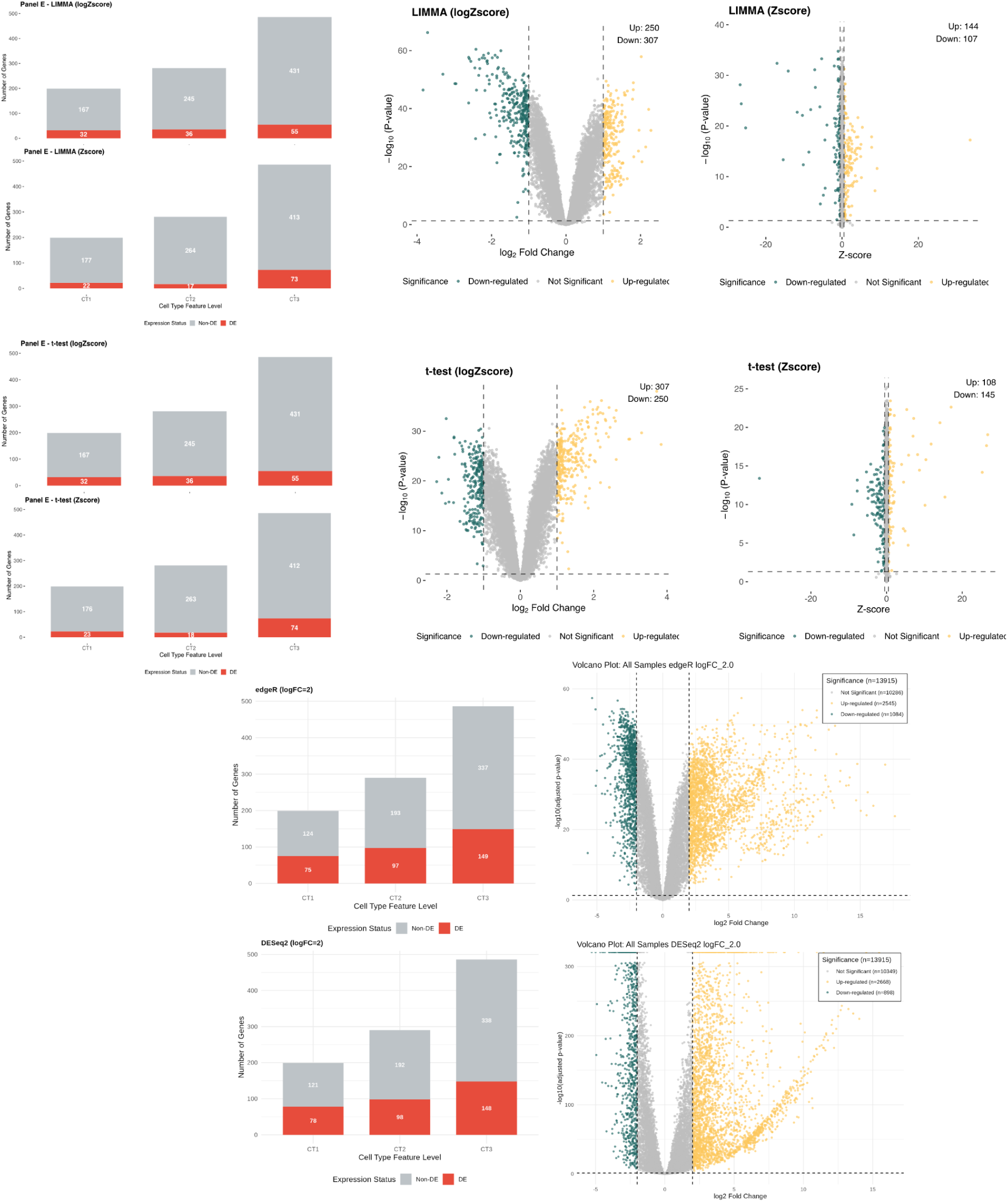
Differential expression analysis using multiple statistical methods. Four statistical approaches (LIMMA, paired *t*-test, edgeR, and DESeq2) were applied to identify differentially expressed genes across three cell type levels. For each method, bar charts show the distribution of DE (red) and non-DE (gray) genes at each level, while volcano plots visualize gene significance and fold changes. The number of DE genes increases with higher cell-type feature levels across all methods, with edgeR and DESeq2 detecting substantially more DE genes than LIMMA and paired *t*-test approaches.

**Figure S3:**
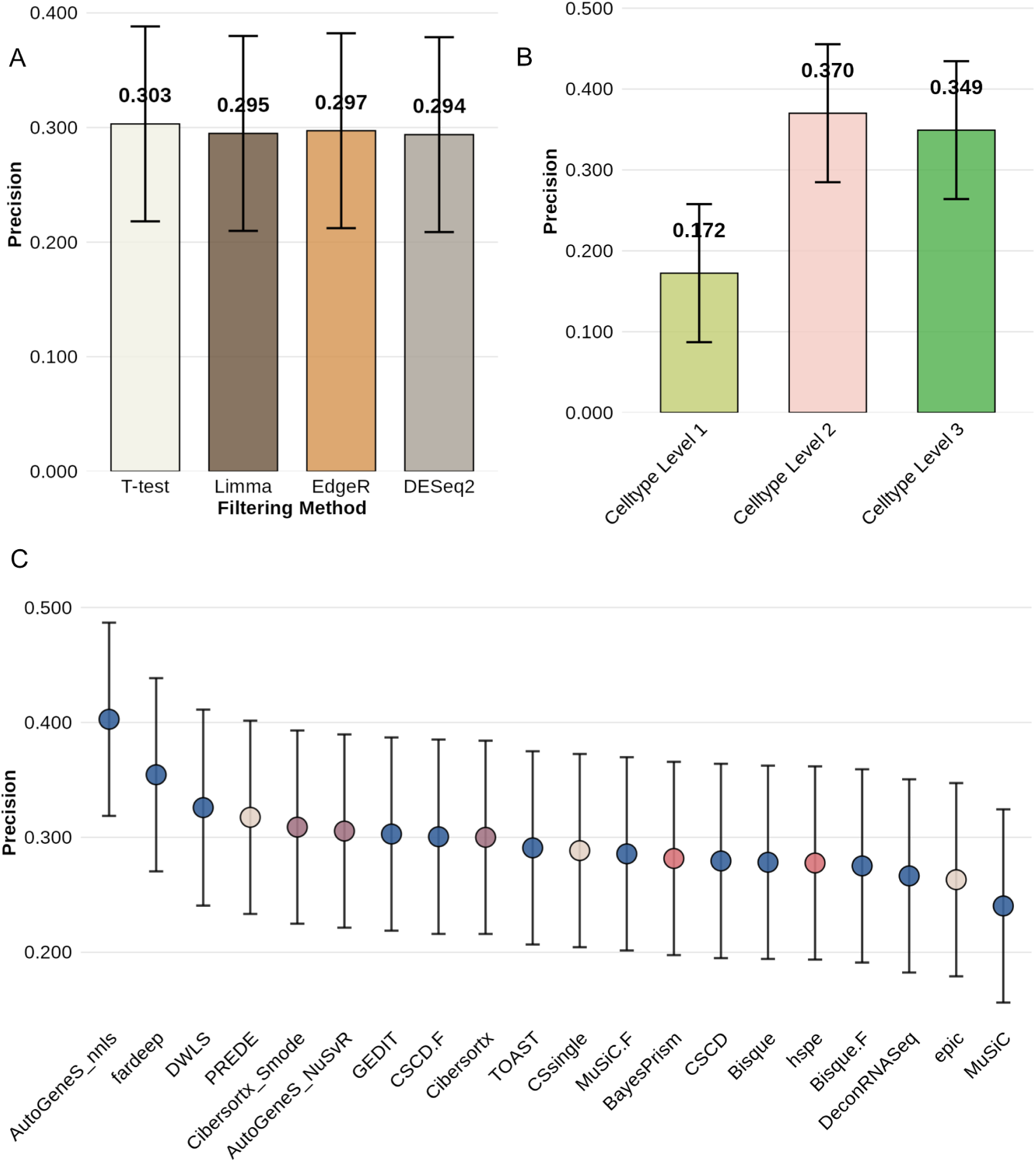
Clinical precision analysis across filtering methods, cell type levels, and deconvolution algorithms. **(A)** Comparison of precision across four filtering methods (Paired *t*-test, Limma, EdgeR, DESeq2) shows similar performance, with mean precision values ranging from 0.294 to 0.303. **(B)** Precision stratified by cell type levels demonstrates higher performance at Celltype Level 2 (0.370) and Level 3 (0.349) compared to Level 1 (0.172), indicating improved clinical prediction accuracy with more granular cell type resolution. **(C)** Precision across 15 deconvolution algorithms reveals AutoGeneS_mls, fardeep, and DWLS as top performers (precision > 0.3), while MuSiC shows the lowest precision.

**Figure S4:**
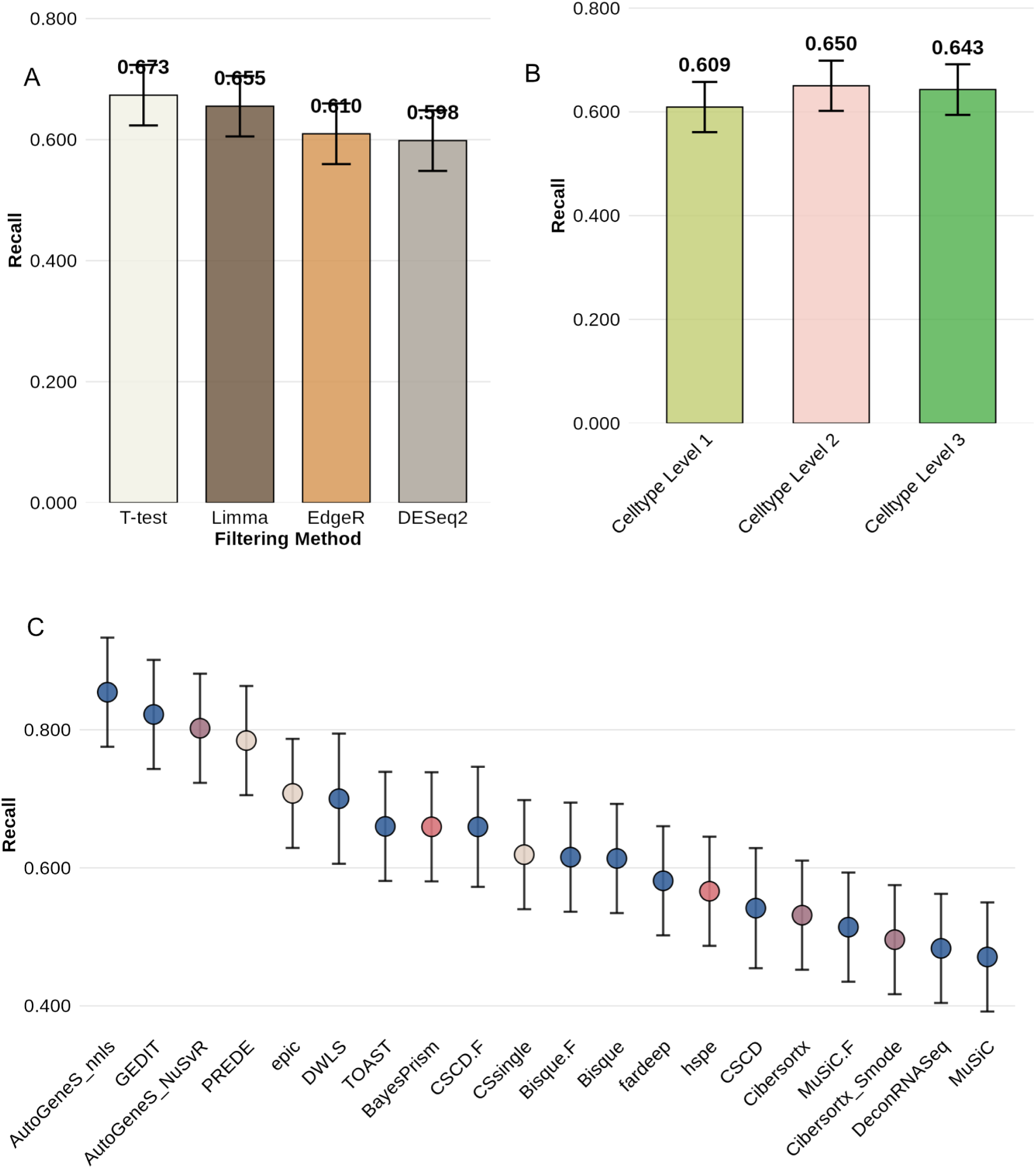
Clinical recall analysis across filtering methods, cell type levels, and deconvolution algorithms. **(A)** Comparison of recall across four filtering methods shows Paired *t*-test achieved the highest performance (0.673), followed by Limma (0.655), EdgeR (0.610), and DESeq2 (0.598). **(B)** Recall stratified by cell type levels demonstrates consistent performance across all three levels, with Level 2 (0.650) and Level 3 (0.643) showing slightly higher recall than Level 1 (0.609), indicating that more detailed cell type resolution maintains sensitivity in clinical prediction. **(C)** Recall across 15 deconvolution algorithms reveals AutoGeneS, and GEDIT as top performers (recall *>* 0.80), while MuSiC, DeconRNASeq, and Cibersortx show lower recall.

**Figure S5:**
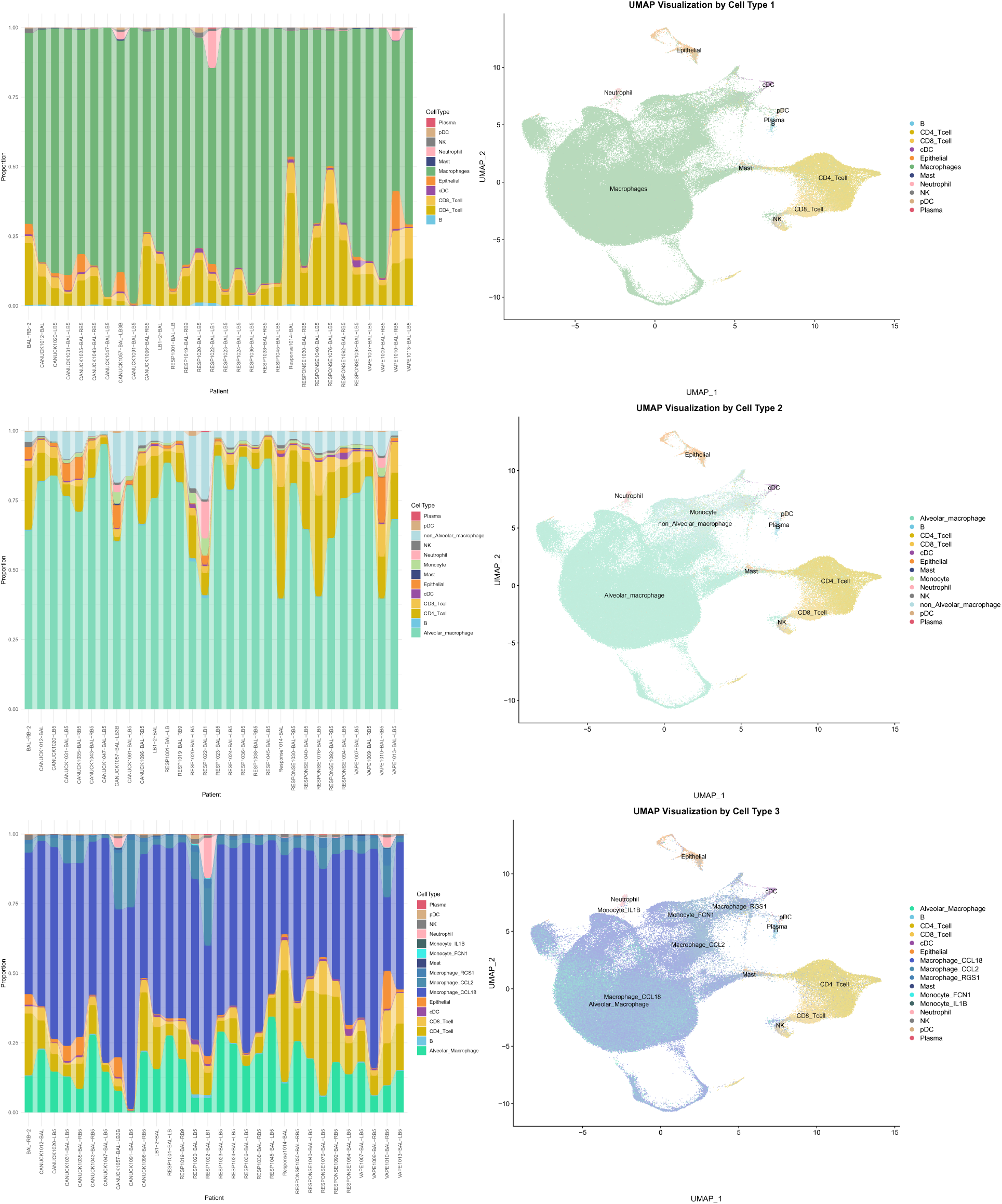
ScRNA-seq cell type composition and UMAP visualization at three annotation levels. The figure visualizes cell type proportions per sample (left) and UMAP projections (right) at three hierarchical cell type annotation levels. **Cell Type Level 1 (top):** Broad classification reveals 11 major populations. **Cell Type Level 2 (middle):** Intermediate resolution identifies 13 cell types. **Cell Type Level 3 (bottom):** High-resolution annotation reveals 18 distinct cell states.

